# Equations governing dynamics of excitation and inhibition in the mouse corticothalamic network

**DOI:** 10.1101/2020.06.03.132688

**Authors:** I-Chun Lin, Michael Okun, Matteo Carandini, Kenneth D. Harris

## Abstract

Although cortical circuits are complex and interconnected with the rest of the brain, their macroscopic dynamics are often approximated by modeling the averaged activities of excitatory and inhibitory cortical neurons, without interactions with other brain circuits. To verify the validity of such mean-field models, we optogenetically stimulated populations of excitatory and parvalbumin-expressing inhibitory neurons in awake mouse visual cortex, while recording population activity in cortex and in its thalamic correspondent, the lateral geniculate nucleus. The cortical responses to brief test pulses could not be explained by a mean-field model including only cortical excitatory and inhibitory populations. However, these responses could be predicted by extending the model to include thalamic interactions that cause net cortical suppression following activation of cortical excitatory neurons. We conclude that mean-field models can accurately summarize cortical dynamics, but only when the cortex is considered as part of a dynamic corticothalamic network.

## Introduction

Brain activity arises from the interactions of large neuronal populations via complex synaptic circuits within and across brain regions. Can the dynamics of such a complex system be accurately summarized by simple quantitative laws? Such laws exist in other fields of science: statistical mechanics shows that physical systems, despite having vast numbers of microscopic, chaotic degrees of freedom, can exhibit a small number of macroscopic degrees of freedom obeying simple quantitative laws to high accuracy.

Multiple efforts have sought to identify simple quantitative laws for the dynamics of cortical circuits. In a landmark study, Wilson and Cowan (Wilson and Cowan, 1972, 1973) proposed equations that could govern the dynamics of circuits comprised of excitatory and inhibitory neurons. The Wilson-Cowan (WC) equations describe only two degrees of freedom – the total summed activities of excitatory and inhibitory neuronal populations – instead of a high-dimensional system describing the precise combination of excitatory and inhibitory neurons active at any moment. This approach, known as mean-field dynamics, has led to substantial theoretical developments (Buice and Chow, 2013; Hertz et al., 2004; Renart et al., 2010; Renart et al., 2007; van Vreeswijk and Sompolinsky, 1996) and has been extended to broader circuits including other brain structures such as the thalamus (Breakspear, 2017; Victor et al., 2011). The WC equations have had a major impact on experimental neuroscience, promoting the idea that to understand cortical functions one should measure the activities of excitatory and inhibitory neurons and characterize their interactions (Anderson et al., 2000; Atallah et al., 2012; Ben-Yishai et al., 1995; Haider et al., 2006; Isaacson and Scanziani, 2011; Kerlin et al., 2010; Lee et al., 2012; Mann et al., 2009; Monier et al., 2003; Murphy and Miller, 2009; Okun and Lampl, 2008; Ozeki et al., 2009; Somers et al., 1995).

Despite their importance and impact, the WC equations have seen little experimental testing. Mean-field dynamical-system models have been shown to produce oscillations that resemble those seen in neural circuits (Akam et al., 2012; Chaudhuri et al., 2015; Murphy and Miller, 2009; Ozeki et al., 2009; Rubin et al., 2015; Tsodyks et al., 1997). Yet, the fact that a model reproduces some features resembling neural activity does not mean that it captures the underlying dynamics. A stronger test is to verify whether the model can accurately predict the effect of experimental perturbations to its components.

Here, we test the ability of the WC equations, and extensions thereof, to quantitatively capture the macroscopic dynamics of cortical circuits. We recorded from the primary visual cortex (V1) of quietly awake mice and estimated the averaged population activity of putative excitatory and fast-spiking inhibitory neurons. We delivered brief test pulses using dichromatic optogenetics to selectively stimulate excitatory and/or parvalbumin-expressing inhibitory populations in V1 with single or paired pulses. The responses of V1 to paired pulses showed a reliable but unexpected interaction, which could not be captured by a purely cortical network model. Additional recordings in the lateral geniculate nucleus (LGN) suggested that the thalamus shapes the response to the activation of the cortical excitatory, but not inhibitory, population. Furthermore, a mean-field model of the thalamocortical dynamics could predict quantitatively responses to all stimuli. We conclude that macroscopic cortical dynamics can be well described by mean-field models, but only if the cortex is considered as part of a broader system that includes connections to and from the thalamus.

## Results

To study and model the dynamics of cortical excitation and inhibition, we recorded population activity from V1 of quietly awake mice, classifying spikes from putative excitatory (E) and fast-spiking inhibitory (I) neurons by their width. We probed their dynamics by optogenetically manipulating these neuronal populations with brief laser pulses in mice expressing different opsins in E and I neurons. We first describe experiments that validate this approach.

### Responses of single cortical neurons to an optogenetic excitatory pulse

To investigate the effect of excitatory optogenetic stimulation on individual excitatory neurons, we used juxtacellular recordings. The experiments were conducted in quietly awake mice of Thy1-ChR2-YFP line 18 (Thy18, expressing ChR2 in layer V pyramidal neurons; Arenkiel et al. (2007)); a single pulse of 1-, 2, or 4-ms duration from a 445 nm laser at 100 mW power was delivered to the surface of the brain every 1.6 to 3 s to stimulate the cortical E population (an “E pulse”; **Fig. 1A,B**). We recorded juxtacellularly from individual V1 neurons (50 neurons in 4 mice; these were primarily E neurons based on their spike waveforms, **Fig. 1D**, inset). An E pulse triggered short bursts of up to 5 spikes within 20 ms from the pulse: 30 out of 50 neurons showed a strong initial response (average ≥1 spike within 20 ms from the pulse; **Fig. 1C,D** and **Supplementary Fig. S1A**).

**Fig. 1.**
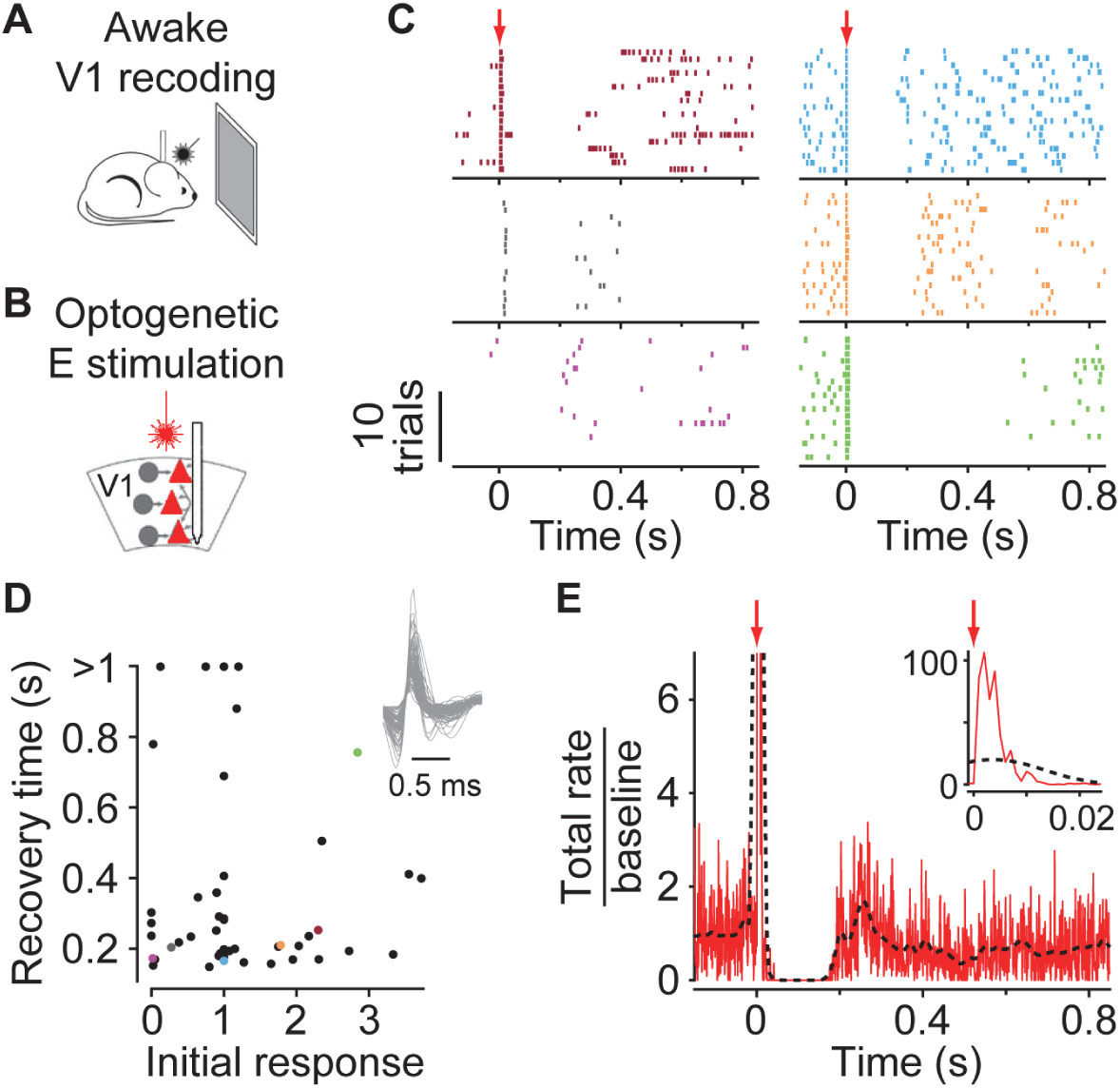
Responses of single cortical neurons to an optogenetic excitatory pulse. (A) Single neurons were recorded juxtacellularly in V1 of quietly awake, head-restrained mice of Thy18 strain, in front of blank monitors. (B) The cortical E population was optogenetically stimulated by a brief pulse from a 445 nm laser (an ‘E pulse’, red color indicates the stimulation of E neurons, rather than the laser wavelength). (C) Raster plots showing responses of six putative E neurons to a 2-ms E pulse. (D) Summary of all recorded neurons’ responses to a 2-ms E pulse. *X*-axis: initial response, mean number of spikes fired within 20 ms from the pulse; *y*-axis: recovery time, time when activity recovered to 50% baseline. Colored dots correspond to neurons in (C). Inset: mean waveforms of all recorded neurons. *n* = 50 neurons, 4 mice. (E) Peristimulus time histogram of responses from a virtual population, generated by summing trial-averaged responses of all recorded neurons then normalizing the summed response to have baseline rate = 1. Dashed black line: Response smoothed by a 40-ms Hamming window. Inset: expanded time scale.

Strikingly, all recorded neurons, regardless of whether they showed an initial response to an E pulse, were suppressed for more than 150 ms post-stimulation (**Fig. 1C–E)**. This suppression was evident in the spike rasters (**Fig. 1C**), and its duration did not depend on the magnitude of the initial response (**Fig. 1D**; Spearman’s rank correlation coefficient: −0.02, *p* = 0.90). This suppression resembled the prolonged suppression previously observed in response to electrical brain stimulation as well as flash stimuli (Douglas and Martin, 1991; Dreifuss et al., 1969; Funayama et al., 2016; Pei et al., 1991). To estimate the population dynamics evoked by an E pulse based on the juxtacellular data, we pooled all recorded neurons to create a virtual population and estimated its rate by summing the trial-averaged responses of all neurons. The virtual-population rate confirmed the impression given by single-cell rasters: delivering a laser pulse to stimulate the E population activated the E neurons, followed by suppression that lasted roughly 150 ms and a rebound to approximately twice the baseline activity (**Fig. 1E**). In individual neurons, this rebound did not correlate with the magnitude of the initial response (Spearman’s rank correlation coefficient: 0.18, *p* = 0.20). The initial response was transient yet very strong, rising to over 100 times the baseline rate when smoothed at 1 ms (**Fig. 1E**, inset). This sharp peak reflected the very precisely timed response of E neurons to optogenetic stimulation, signifying highly synchronous population activity (**Supplementary Fig. S1**). Even when smoothed at 40 ms (necessary for direct comparison to extracellular recordings later), the peak rate of this initial response was still 20 times the baseline rate.

### Population responses to single optogenetic excitatory or inhibitory pulse

To measure and manipulate the activity of cortical E and I populations simultaneously, we used multi-site silicon probes with targeted dichromatic optogenetics (*n* = 19 mice, **Supplementary Tables S1** and **S2**). To distinguish the firing of the two populations, we took advantage of the narrow spike waveform of fast-spiking inhibitory cells (Bartho et al., 2004) and developed a “clusterless” method to estimate the total rates of wide and narrow spikes detected by the silicon probe (Supplemental Information S1.6 and **Supplementary Fig. S2**). To verify that our methods for stimulating and recording selected neuronal populations worked as intended, we studied the responses to single pulses of either wavelength: by expressing ChR2 opsin in pyramidal neurons (E) and C1V1_T/T_ opsin (Yizhar et al., 2011) in parvalbumin-expressing cells (I) of V1, we could selectively drive the two populations with pulses of different wavelengths – 445 nm for E cells; 561 nm for I cells.

Stimulating E and I populations with optogenetic pulses resulted in different impulse-response dynamics. Consistent with the juxtacellular data, delivering an optogenetic pulse to stimulate E neurons (an “E pulse”; 445 nm in either Thy18 mice or PV^cre^;Thy18 mice injected with C1V1_T/T_ virus) elicited an immediate response in both E and I populations (the latter as expected via monosynaptic drive to I neurons), followed by sustained suppression (∼150 ms) and then a rebound in activity in both populations (**Fig. 2A**). By contrast, delivering an optogenetic pulse to stimulate I neurons (an “I pulse”; 445 nm in PV^cre^;Ai32 mice and 561 nm in PV^cre^;Thy18 mice injected with C1V1_T/T_ virus) caused an immediate response in the I, but not the E, population (**Fig. 2B**). Following this immediate response, the I pulse evoked in both populations prolonged suppression, which was nevertheless shorter and less pronounced than the one evoked by an E pulse and not followed by a strong rebound (**Fig. 2B**). The time course of the E population response to an E pulse closely matched the one obtained from juxtacellular recordings, except for the first 20 ms after the pulse, during which the extracellular signals severely underestimated the highly synchronous activity seen in juxtacellular recordings (**Fig. 2A**, top). Examining raw traces from extracellular recordings confirmed that there was strong superposition of multiple spikes at this time, together with strong high-frequency fluctuations in field potential, explaining the lower rates of detected spikes (**Supplementary Fig. S1B**).

**Fig. 2.**
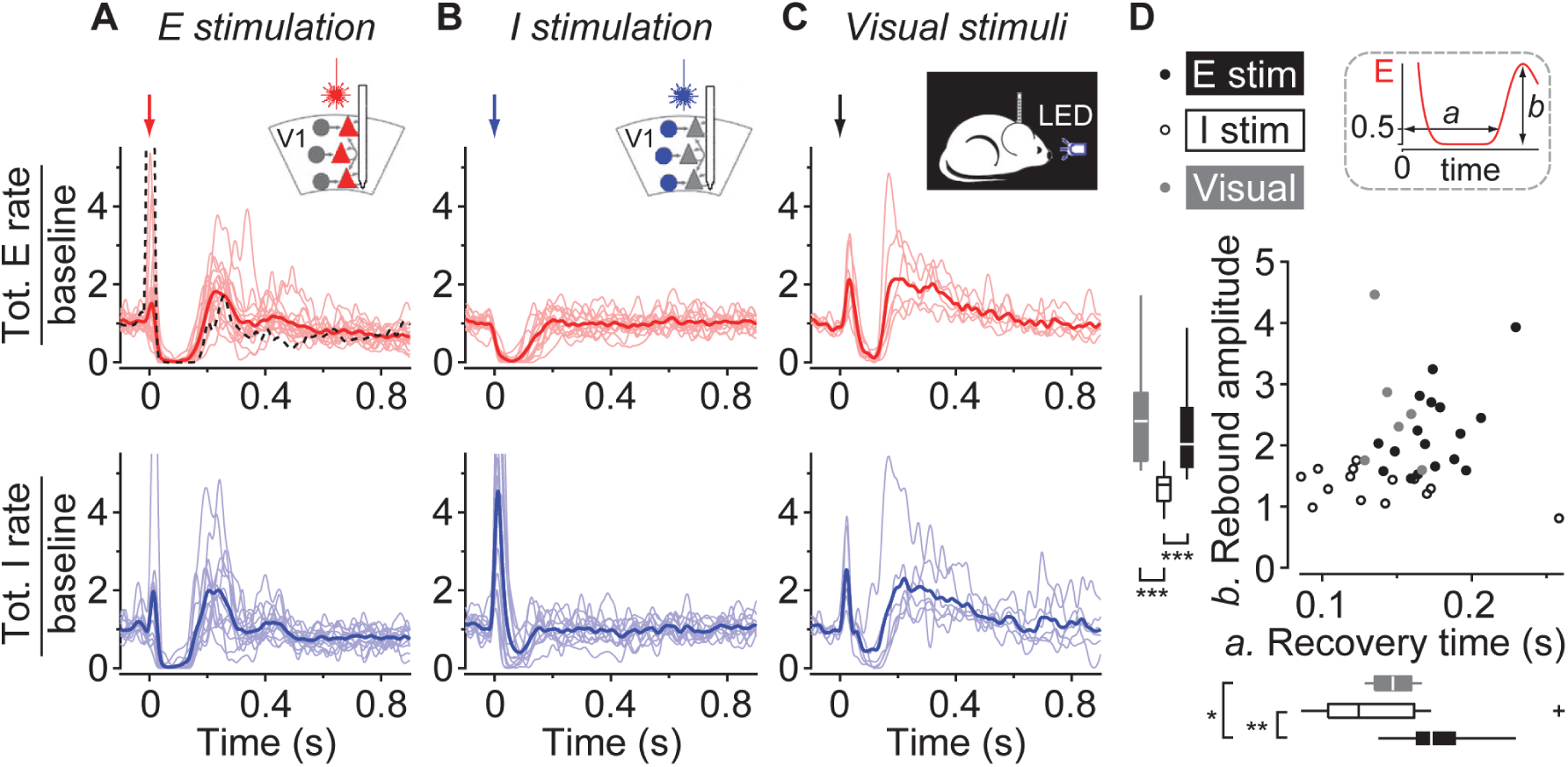
Population responses to an optogenetic excitatory or inhibitory pulse, or to visual stimuli. (A) Peristimulus time histograms of cortical E (red, top) and I (blue, bottom) population responses to an E pulse (marked by an arrow). Thin red/blue lines: data to one pulse duration from individual recording sessions; bold red/blue line: session average. Dashed black line: the virtual-population rate from juxtacellular recordings as in Fig. 1E. Note that E population responses recorded extracellularly and juxtacellularly match closely except for the initial response. *n* = 17 sessions, 17 mice; I population responses from two sessions were omitted because of low baseline rates (<15 spikes/s). (B) As in (A) but for responses to an I pulse. *n* = 14 sessions, 13 mice. I population responses from two sessions were omitted because of low baseline rates (<15 spikes/s). (C) As in (A) but for responses to a brief light flash to the contralateral eye of a mouse in darkness. *n* = 6 sessions, 5 mice. (D) Recovery time and rebound amplitudes of E population responses, across sessions. Each symbol (● E stimulation; ○ I stimulation; ● visual stimuli) denotes data averaged across all pulse durations in one session. Box plots show median as bar, 25–75% interval as box, and 1.5 IQR as whiskers. Mixed model ANOVA: *, *p* < 0.05; **, *p* < 0.01; ***, *p* < 0.001. Inset: recovery time is the time when the E population response reaches 50% of its baseline activity; rebound amplitude is the peak E population response post-suppression.

The complete and long-lasting suppression of cortical activity by an E pulse was not specific to optogenetic stimulation and could also be evoked by flash visual stimuli (**Fig. 2C**). To show this, we carried out additional recordings in five wild-type mice, while briefly flashing an LED to generate a pulsatile visual stimulus. The evoked dynamics were similar to those following an E pulse: initial firing, prolonged suppression and a strong rebound in activity (**Fig. 2C**).

An E pulse and an I pulse evoked significantly different population dynamics, with the E pulse triggering longer suppression and stronger rebounds than the I pulse (**Fig. 2D**). We characterized the evoked dynamics by two numbers: the recovery time, defined as the time the E population response took to recover to 50% of its baseline activity, and the rebound amplitude, defined as the peak E population response post-suppression (**Fig. 2D**, inset). We pooled data from all pulse durations within a recording session, as we did not observe a major effect of pulse duration on the evoked dynamics (**Supplementary Fig. S3**; E pulse: *p* = 0.10, 34 data points in 17 mice; I pulse: *p* = 0.17, 26 data points in 13 mice; mixed model ANOVA). Compared to an I pulse, an E pulse caused longer suppression (E pulse: 174 ± 6ms, mean ± SE, 17 sessions in 17 mice; I pulse: 137 ± 12 ms, 14 sessions in 13 mice; *p* = 10^−3^, mixed model ANOVA) and larger rebound amplitudes (E pulse: 2.22 ± 0.16, mean ± SE times baseline; I pulse: 1.33 ± 0.07; *p* = 5×10^−5^, mixed model ANOVA). An LED flash to the contralateral eye (flash visual stimulus) triggered intermediate-length suppression (147 ± 6 ms, mean ± SE, 6 sessions in 5 mice. E pulse vs. flash: *p* = 0.01; I pulse vs. flash: *p* = 0.58, mixed model ANOVA) and a strong rebound (2.58 ± 0.42, mean ± SE times baseline. E pulse vs. flash: *p* = 0.28; I pulse vs. flash: *p* = 2×10^−4^, mixed model ANOVA).

### Population responses to paired optogenetic pulses

To further constrain our models, we recorded responses to pairs of E and I pulses (“paired pulse”) presented in different orders at several interpulse intervals (IPIs). The data were collected from nine PV^cre^;Thy18 mice injected with C1V1_T/T_ virus, and we delivered E and I pulses with 445- and 561-nm lasers (**Supplementary Tables S2** and **S3**).

The paired-pulse experiments gave an unexpected result: regardless of whether the E pulse preceded or followed the I pulse, the suppression tended to end at a constant time after the E pulse (**Fig. 3**). When an I pulse preceded an E pulse (I→E stimulation), the evoked suppression ended at a fixed time after the second pulse: the recovery time increased from 181 to 261 ms as the IPI increased from 12 to 90 ms (**Fig. 3A**; recovery time vs. IPI: Spearman’s rank correlation coefficient 1, *p* = 3×10^−3^). By contrast, when an E pulse preceded an I pulse (E→I stimulation), the suppression ended at a fixed time after the first pulse: as IPI increased from 12 to 90 ms, the recovery time only increased moderately from 159 to 175 ms (**Fig. 3B**; recovery time vs. IPI: Spearman’s rank correlation coefficient 0.77, *p* = 0.10). This relationship became clearer when superimposing the mean responses at different IPIs (**Fig. 3C–F**). To further verify, we compared the slopes of the linear fits to recovery time as a function of IPI (over a range of 5 to 100 ms) for I→E and E→I stimulation (**Fig. 3G**). For I→E stimulation, the slope was close to 1, whereas for E→I stimulation, the slope was close to 0, indicating that in both cases the suppression ended at a fixed time after an E pulse. While the values of the slopes varied from session to session, the slopes of I→E and E→I stimulation were significantly different within each session (**Fig. 3H**. Median slopes of I→E and E→I stimulation across sessions: 1.0 and 0.2, *p* = 4×10^−3^; Wilcoxon signed-rank test, 9 sessions in 9 mice). We also applied two consecutive E pulses and I pulses (E→E and I→I stimulation; **Supplementary Fig. S4**). The slopes of E→E and I→I stimulation varied greatly from session to session but did not significantly differ from each other (*p* = 0.57; Wilcoxon signed-rank test, 9 sessions in 9 mice). Both slopes were significantly larger than the slopes of E→I stimulation (*p* = 0.02 and 4×10^−3^ for E→E and I→I stimulation; Wilcoxon signed-rank test, 9 sessions in 9 mice), but smaller than the slopes of I→E stimulation (*p* = 4×10^−3^ for both).

**Fig. 3.**
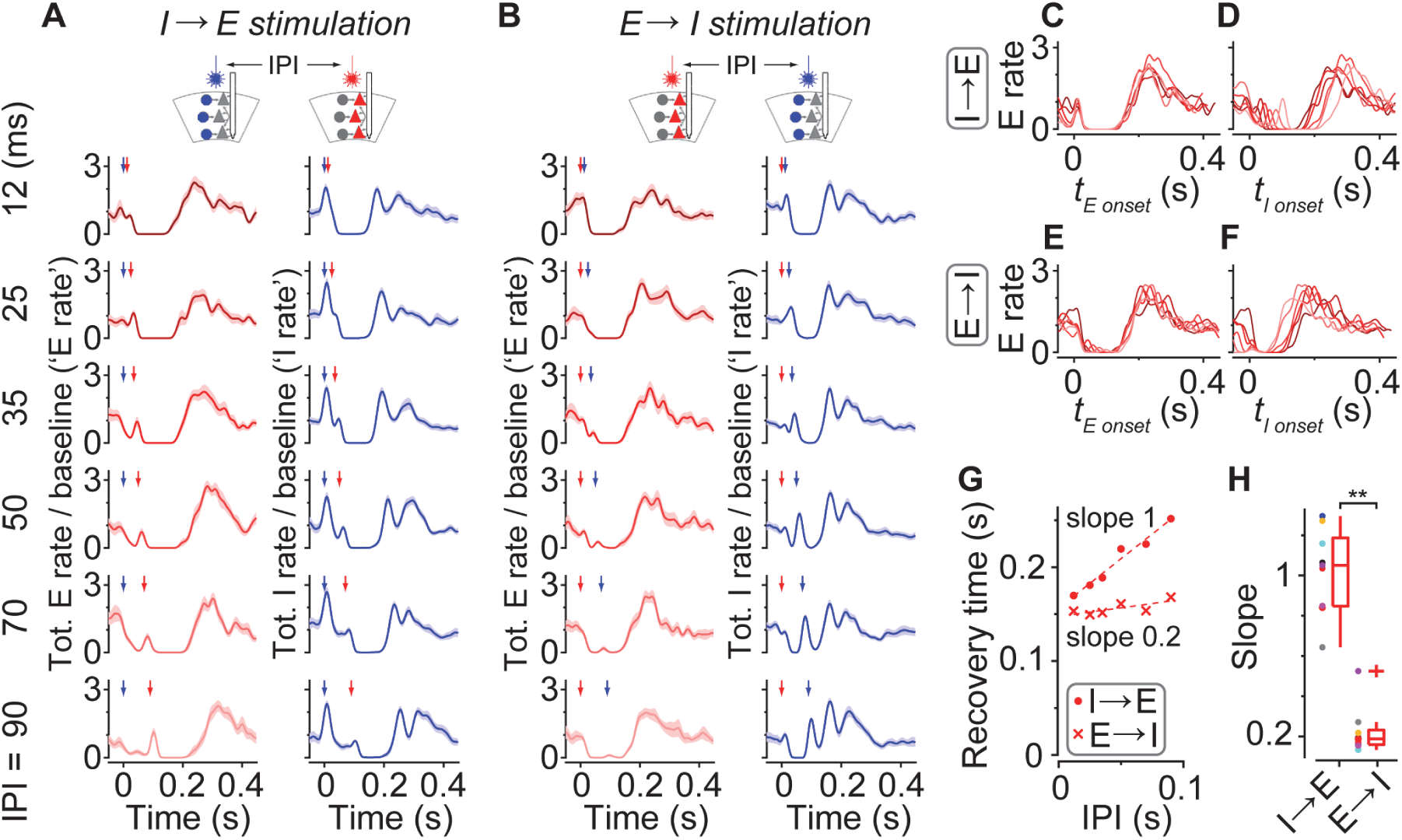
Population responses to paired optogenetic pulses. (A) Peristimulus time histograms of cortical E and I population responses to optogenetic stimulation of first I then E neurons (I→E stimulation) at various interpulse intervals (IPIs), for an example session. Red trace (left column): mean E population rate; Blue trace (right column): mean I population rate. Each row shows the mean response across trials with a fixed IPI, normalized so baseline rate is 1. Red and blue arrows: E and I pulses. Shaded area: s.e.m. across trials in a session. (B) As in (A) but for responses to optogenetic stimulation of first E then I neurons (E→I stimulation). (C) E population responses to I→E stimulation for the same session, overlaid and re-aligned to the E pulse. Colors represent the IPIs, matching rows in (A). (D) As in (C) but aligned to the I pulse. (E,F) As in (C,D) but for E→I stimulation. Colors match rows in (B). (G) Recovery time of E population responses to I→E (●) and E→I (×) stimulation as a function of IPI. Dashed lines, linear fits. (H) Box plots summarizing the slopes of the linear fits to recovery time vs. IPI up to 100 ms across 9 sessions in 9 mice. Bar, median; box, 25–75% interval; whiskers, 1.5 IQR. Each dot marks data from a session; data from the same session share the same color. Wilcoxon signed-rank test: **, *p* < 0.01.

### The WC model cannot account for multiple features of the impulse-response dynamics

We next sought a simple set of equations that could predict the responses of cortical E and I populations to these test pulses. We started with the WC equations that describe the dynamics of two populations of neurons, one excitatory and one inhibitory, each making a recurrent connection onto itself and a reciprocal connection onto each other (**Fig. 4A**). Summarizing the total firing of the E population as *E*(*t*) and the total firing of the I population as *I*(*t*), the equations are:

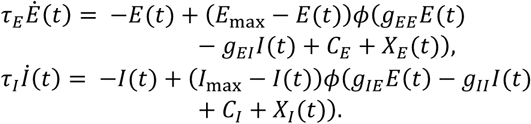

**Fig. 4.**
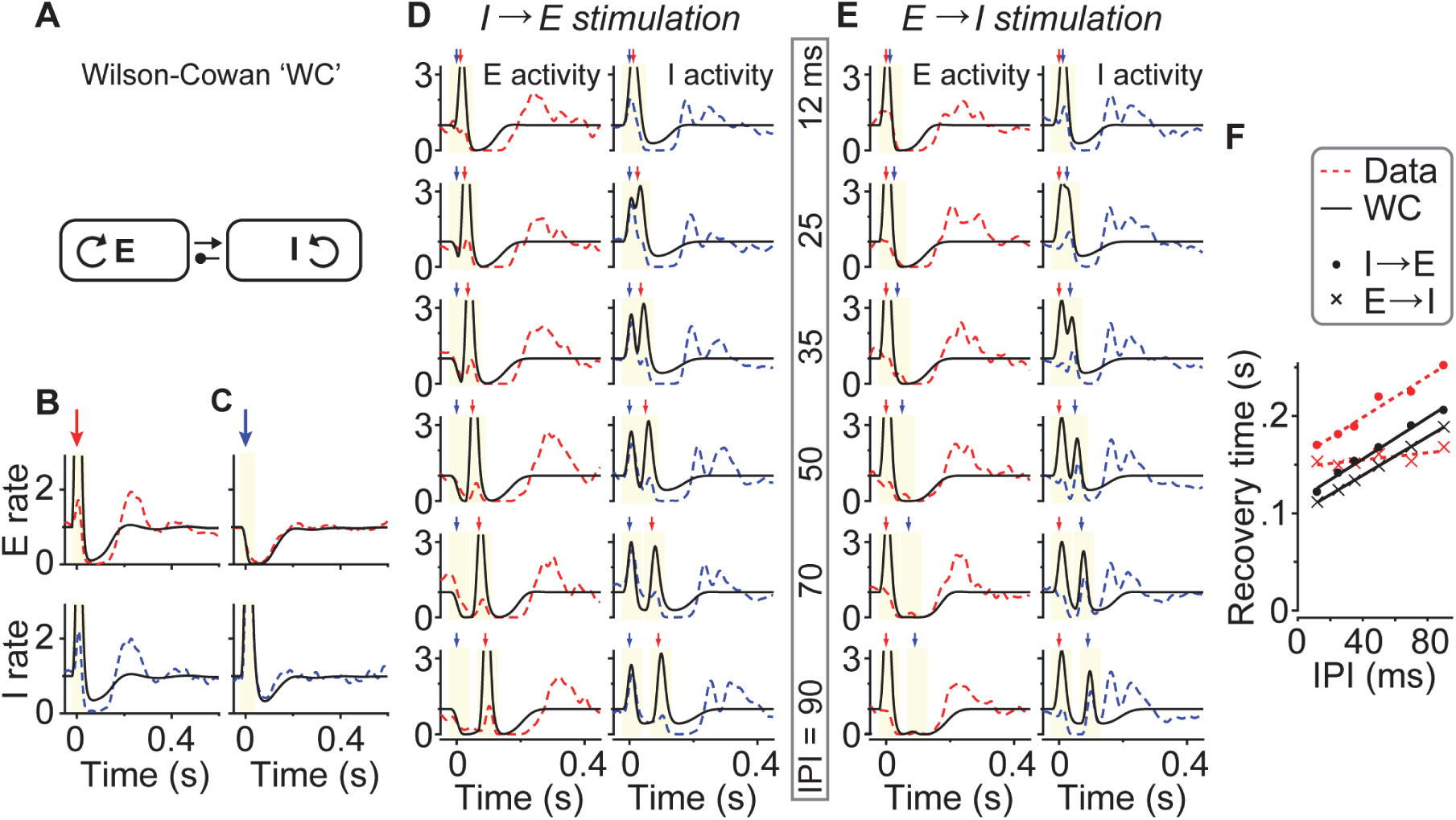
The Wilson-Cowan equations could not model the observed cortical dynamics. (A) A schematic of the Wilson-Cowan (WC) equations. Two populations of neurons, one excitatory and one inhibitory, are recurrently and reciprocally connected. Each population receives a constant input *C* and an optogenetic-pulse input *X* (not shown). (B,C) Best fit of the WC model to single optogenetic-pulse experimental data. Peristimulus time histograms of experimentally measured E (red dashed, top) and I (blue dashed, bottom) population responses to single E (B) and I (C) pulses. Solid black lines: WC model predictions. Arrows: optogenetic pulses. Ivory regions: periods excluded from the loss function. Experimental data shown here were averaged across 11 sessions in 11 mice; model predictions were averaged across all folds of cross-validation and all sessions. (D,E) Best fit of the WC model to paired-pulse experimental data, for an example session. Red and blue dashed lines: measured E and I population responses, replotted from Fig. 3A,B. Black solid lines: WC model predictions. Red and blue arrows: E and I pulses. (F) Recovery time of experimentally measured (red) and WC-predicted (black) E population responses to I→E (●) and E→I (×) stimulation as a function of IPI. Red dashed and black solid lines are linear fits to experimental data and WC model predictions.

Here, *E*(*t*) and *I*(*t*) denote the level of activity in each neuronal population, whose baseline rate is scaled to 1. *X*(*t*) and *C* represent external inputs to the populations: the former represents the transient effect of an optogenetic pulse; the latter represents a constant input. We took the activation function *ϕ*(*x*) to be a rectified linear function, *ϕ*(*x*) = max(*x*, 0). The parameters *g*_*EE*_, *g*_*IE*_, *g*_*EI*_, and *g*_*II*_ represent the strengths of recurrent and reciprocal connections within and between the two populations. *τ*_*E*_ and *τ*_*I*_ denote the time constants of the E and I populations; *E*_max_ and *I*_max_ represent the maximum possible activation level of the populations.

We fit the model parameters by minimizing a loss function which quantifies the difference between the dynamics predicted by the model and the firing of E and I populations (Supplemental Information S1.8 and S1.9). This loss function did not penalize over-prediction of E and I population responses immediately after optogenetic stimulation, because our extracellular measurements underestimate the true firing rates (**Fig. 2A**, top, and **Supplementary Fig. S1**). We evaluated the model’s performance using cross-validation (Supplemental Information S1.10), searching for optimal parameters on the averaged response over one subset of trials, and evaluating the model’s performance on the averaged response over held-out trials for single-pulse data (**Supplementary Fig. S5A**). For both the visual-stimulus data and the paired-pulse data, we used an out-of-sample approach, in which the model was trained using data from LED flashes of different durations (visual-stimulus data) or paired pulses of different IPIs (paired-pulse data) to the test set, ensuring that the model could generalize across stimulus conditions (**Supplementary Fig. S5B,C**). The WC model could not describe all aspects of the cortical dynamics evoked by optogenetic pulses (**Fig. 4B,C**) and flash visual stimuli. For responses to an E pulse, even with optimal parameters the WC model could not correctly predict the rebound amplitudes (measured and WC-predicted rebound amplitudes across sessions: medians = 2.20 and 1.02 times baseline, *p* = 9.8×10^−4^; Wilcoxon signed-rank test, 11 sessions in 11 mice), the suppression duration (data and WC predictions across sessions: medians = 164 and 139 ms, *p* = 9.8×10^−4^; Wilcoxon signed-rank test, 11 sessions in 11 mice), nor that these suppression durations were reliably longer than the ones evoked by I pulses (data, *p* = 6.8×10^−3^; WC predictions: *p* = 1; Wilcoxon signed-rank test on durations of suppression evoked by E vs. I pulses, 11 sessions in 11 mice). The model also could not predict the near-complete suppression in the I population triggered by an E pulse (measured and WC-predicted minimum I activities *I*_min_ across sessions: medians = 0 and 0.34 times baseline, *p* = 9.8×10^−4^; Wilcoxon signed-rank test, 11 sessions in 11 mice). Furthermore, the model was unable to generate strong enough rebounds after flash visual stimuli (measured and WC-predicted response characteristics across sessions: recovery time, medians = 146 and 142 ms, *p* = 0.17; rebound amplitudes, medians = 2.39 and 1.59 times baseline, *p* = 9×10^−5^; *E*_min_, medians = 0.06 and 0 times baseline, *p* = 2×10^−4^; *I*_min_, medians = 0.09 and 0.05 times baseline, *p* = 0.052; Wilcoxon signed-rank test, 4 conditions per session, 6 sessions in 5 mice). These specific failures of the WC model are to be expected in a dynamical system where only activity of I cells can suppress E cells: in such models, the I rate can never reach zero, and E activity must begin increasing at least as soon as I activity has hit its minimum.

The WC model also failed to reproduce the key finding of the paired-pulse experimental data (**Fig. 4D–F**), namely that the suppression terminated at a fixed time following an E pulse, irrespective of the order in which the E and I pulses were delivered. Instead, the time course of the suppression predicted by the WC model always followed the timing of the second pulse, regardless of the pulse order (slopes of WC-predicted recovery time to I→E and E→I stimulations as a function of IPI, across sessions: medians = 1.08 and 1.05, *p* = 0.82; Wilcoxon signed-rank test, 9 sessions in 9 mice). Similar results were found for E→E and I→I stimulation (**Supplementary Fig. S6**). Again, this failure is inevitable in a model where only I activity can suppress E firing: suppression of the E population can only occur through driving the I population to high rates, and the timescale of recovery from this suppression will be set by the same parameter *τ*_I_, regardless of whether the I population was driven directly or via the E population.

While the WC model did not accurately capture the responses to an E pulse or paired pulses, it modeled the responses to an I pulse reasonably well (**Fig. 4c)**, albeit with still underestimated rebound amplitudes (measured and WC-predicted response characteristics across sessions: recovery time, medians = 123 and 128 ms, *p* = 0.21; rebound amplitudes, medians = 1.36 and 1.01 times baseline, *p* = 4.9×10^−3^, *E*_min_, medians = 0.01 and 0 times baseline, *p* = 0.32; *I*_min_, medians = 0.15 and 0.33 times baseline, *p* = 0.52; Wilcoxon signed-rank test, 11 sessions in 11 mice). In particular, the WC model was able to capture the suppression of I cells following the initial strong response to the I pulse. This “paradoxical” suppression of I activity following activation of I cells is a feature of “inhibitory stabilized network” models (ISNs), in which recurrent excitation is so strong that it would lead to instability if it were not compensated by strong inhibition (Litwin-Kumar et al., 2016; Murphy and Miller, 2009; Ozeki et al., 2009; Sadeh et al., 2017; Tsodyks et al., 1997). Examining the parameters of the model fits confirmed that experiments with stronger I suppression following an I pulse were fit by models with stronger recurrent excitation (Pearson’s linear correlation between *E*_max_*g*_*EE*_ and minimum I activity *I*_min_: −0.66, *p* = 3×10^−5^).

### Adding slow cortical inhibition improves the accuracy of the WC model but only partially

We next asked whether it is possible to overcome the failures of the WC model by adding slow cortical inhibition. Although synaptic GABAergic inhibition typically operates at a timescale of a few milliseconds, GABA inhibition can also have a slow component, for example, by the non-synaptic action of GABA diffused into the extracellular space acting via GABA_B_ or GABA_A_ slow conductance (Farrant and Nusser, 2005; Scanziani, 2000; Simon et al., 2005; Tamas et al., 2003). We reasoned that adding slow GABAergic inhibition to the model could provide “momentum” to cortical inhibition, improving the fit by allowing continued suppression of E cells even when I cells are also suppressed. To achieve this, we extended the WC model by adding a variable *S* that models slow GABAergic inhibition as a leaky integral of I cell activity (the “WCS model”, **Fig. 5A**):

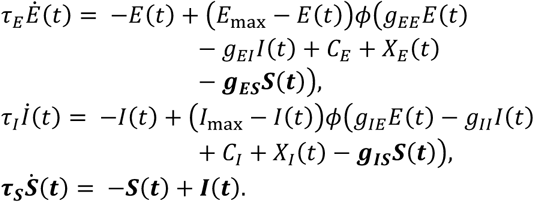

**Fig. 5.**
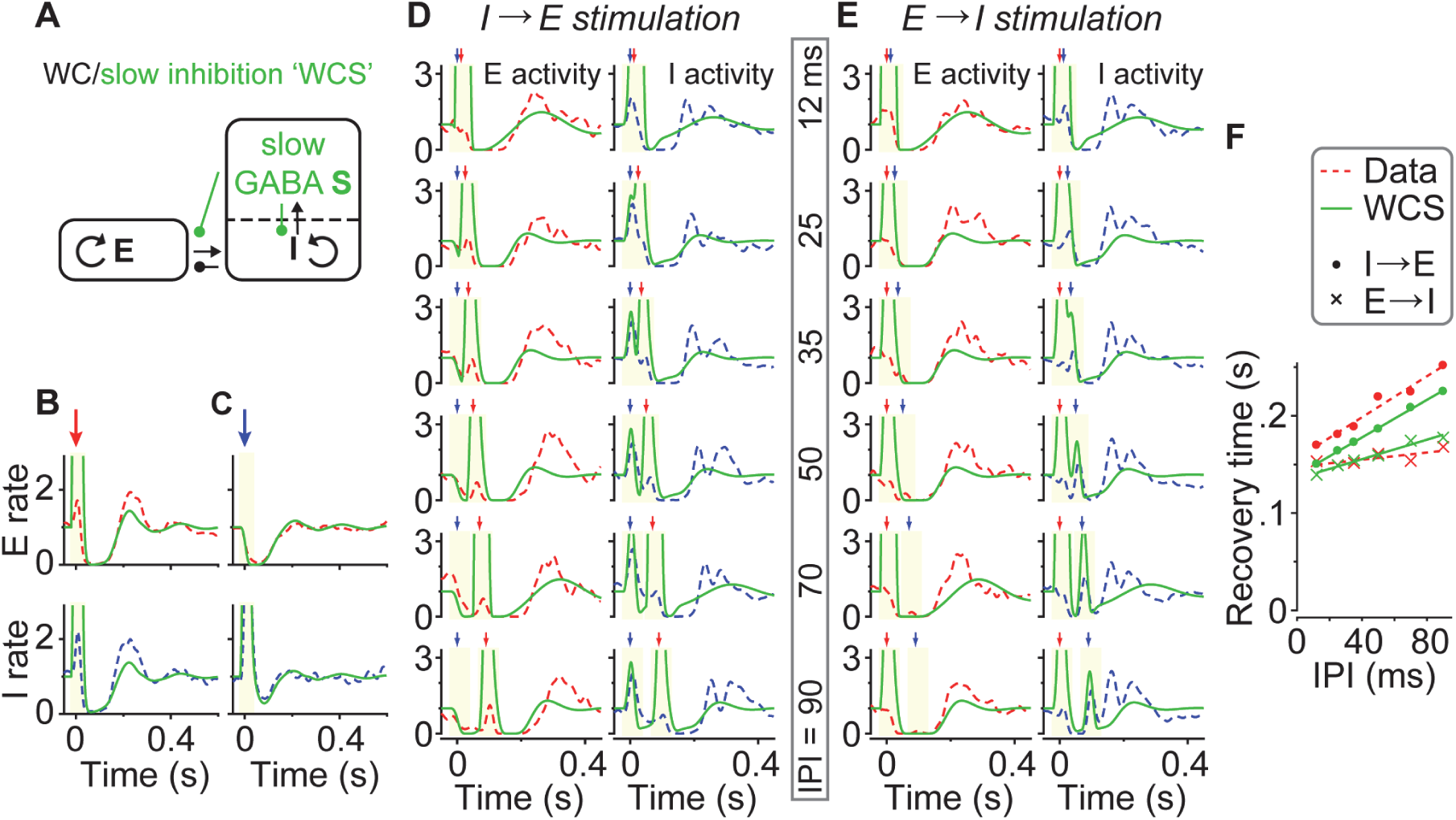
Adding slow cortical inhibition only moderately improved the accuracy of the Wilson-Cowan equations. (A) A schematic of the Wilson-Cowan/slow inhibition (WCS) model. This model contains an additional mechanism modeled after the slow non-synaptic inhibition, allowing for suppression with a longer time constant. (B,C) Best fit of the WCS model to single optogenetic-pulse experimental data. Peristimulus time histograms of experimentally measured E (red dashed, top) and I (blue dashed, bottom) population responses to single E (B) and I (C) pulses. Solid green lines: WCS model predictions. Arrows: optogenetic pulses. Ivory regions: periods excluded in the loss function. Experimental data shown here were averaged across 11 sessions in 11 mice; model predictions were averaged across all folds of cross-validation and all sessions. (D,E) Best fit of the WCS model to paired-pulse experimental data, for an example session. Red and blue dashed lines: measured E and I population responses, replotted from Fig. 3A,B. Green solid lines: WCS model predictions. Red and blue arrows: E and I pulses. (F) Recovery time of experimentally measured (red) and WCS-predicted (green) E population responses to I→E (●) and E→I (×) stimulation as a function of IPI. Red dashed and green solid lines are linear fits to data and WCS model predictions.

In this set of equations, the terms in **bold** are enhancements over the WC model. The time constant of the slow inhibition *τ*_*S*_ (which could be interpreted, for example, as the time course of extracellular GABA clearance) was allowed to differ from *τ*_*I*_; the parameters *g*_*ES*_ and *g*_*IS*_ are the weights of this slow inhibition on the E and I populations. To test this model’s performance, we refit all parameters, using cross-validation as described previously.

Adding the slow-inhibition term moderately improved the model’s accuracy (**Fig. 5B–F**). The WCS model predicted the response to an E pulse better than the WC model, predicting longer suppression following an E pulse (**Fig. 5B;** WCS-predicted suppression durations due to an E pulse and an I pulse across sessions: medians = 159 and 122 ms, *p* = 0.02; Wilcoxon signed-rank test, 11 sessions in 11 mice). Nevertheless, the predicted rebound amplitudes following an E pulse still fell short of the experimental values (measured and WCS-predicted response characteristics across sessions: recovery time, medians = 164 and 159 ms, *p* = 0.21; rebound amplitudes, medians = 2.20 and 1.49 times baseline, *p* = 9.8×10^−4^; *E*_min_, medians = 0 for both, *p* = 0.70; *I*_min_, medians = 0 and 0.03 times baseline, *p* = 0.07; Wilcoxon signed-rank test, 11 sessions in 11 mice). The WCS model accurately modeled responses to an I pulse, particularly the rebound amplitudes (**Fig. 5C**; measured and WCS-predicted response characteristics across sessions: recovery time, medians = 123 and 122 ms, *p* = 0.32; rebound amplitudes, medians = 1.36 and 1.41 times baseline, *p* = 0.17; *E*_min_, medians = 0.01 and 0 times baseline, *p* = 0.12; *I*_min_, medians = 0.15 and 0.12 times baseline, *p* = 0.10; Wilcoxon signed-rank test, 11 sessions in 11 mice). The parameters of this model (32 out of 33 parameter fits) were also consistent with an inhibitory-stabilized regime, as in the WC model.

The WCS-modeled paired-pulse responses were improved over the ones of the WC model (**Fig. 5D–F**). The WCS model came slightly closer to matching the key finding that recovery occurred at a fixed time after the E pulse, whether it came first or second (slopes of WCS-predicted recovery time to I→E and E→I stimulation as a function of IPI, across sessions: medians = 0.98 and 0.96, *p* = 0.01; Wilcoxon signed-rank test, 9 sessions in 9 mice). Nevertheless, the predicted slopes to E→I stimulation were still larger than the experimental values (slopes of measured and WCS-predicted recovery time as a function of IPI across sessions: medians = 0.18 and 0.96, *p* = 4×10^−3^; Wilcoxon signed-rank test, 9 sessions in 9 mice). The model was also unable to describe the dynamics following flash visual stimuli (measured and WCS-predicted response characteristics across sessions: recovery time, medians = 146 and 165 ms, *p* = 7×10^−5^; rebound amplitudes, medians = 2.39 and 1.80 times baseline, *p* = 3×10^−5^; *E*_min_, medians = 0.06 and 0 times baseline, *p* = 4×10^−5^; *I*_min_, medians = 0.09 and 0 times baseline, *p* = 4×10^−4^; Wilcoxon signed-rank test, 4 conditions per session, 6 sessions in 5 mice).

### Geniculate responses to optogenetic stimulation of cortical excitatory and inhibitory neurons

As we were unable to accurately fit our experimental data with purely cortical models, we considered the possibility that cortical dynamics was additionally shaped by interactions with other brain regions. In particular, it has long been established that thalamic inputs exert powerful influence on cortical dynamics in states such as sleep (David et al., 2013; Gent et al., 2018; McCormick and Bal, 1997). Recent work suggests that this is also the case in waking (Poulet et al., 2012) and that cortical activity is substantially reduced when its thalamic input is withdrawn (Guo et al., 2017b; Reinhold et al., 2015; Steriade, 2001). Furthermore, thalamic circuits are known to produce patterns of suppression followed by a rebound (Steriade et al., 1993), similar to what we observed in cortical dynamics following an E pulse. We therefore hypothesized that similar dynamics of suppression and rebound might also be observed in thalamus, in response to optogenetic pulses delivered to the cortex.

To test this hypothesis, we paired optogenetic stimulation of cortical neurons with extracellular recordings in LGN alone, or in LGN and V1 simultaneously. Optogenetically stimulating the cortical E population produced LGN dynamics similar to those seen in V1: initial short-latency activation, followed by prolonged suppression and a rebound in activity (**Fig. 6A**). By contrast, stimulating the cortical I population had only mild effects on LGN, despite noticeable suppression of V1 activity (**Fig. 6B**). These effects were consistent across recording sessions (**Fig. 6C**): an E pulse to the cortex triggered strong, prolonged suppression in LGN (minimum activity post-stimulation: 0.016 ± 0.010, mean ± SE times baseline, not significantly different from 0, one-sample *t*-test; recovery time: 150 ± 12 ms, mean ± SE; 6 recording sessions in 5 mice) and a robust rebound (2.02 ± 0.21, mean ± SE times baseline) in LGN. In addition, the LGN activity recovered before the V1 activity (*p* = 6×10^−17^; mixed model ANOVA, 17 sessions in 17 mice for V1 recordings, 6 sessions in 5 mice for LGN recordings). An I pulse to the cortex, on the other hand, only partially suppressed the LGN activity (minimum activity post-stimulation: 0.344 ± 0.060 mean ± SE times baseline, significantly different from zero, *p* = 4.7×10^−3^; one-sample *t*-test, 6 recording sessions in 5 mice). This suggests that the distinct cortical dynamics in response to an E and an I pulse to the cortex may be due to their fundamentally different consequences on thalamic activity.

**Fig. 6.**
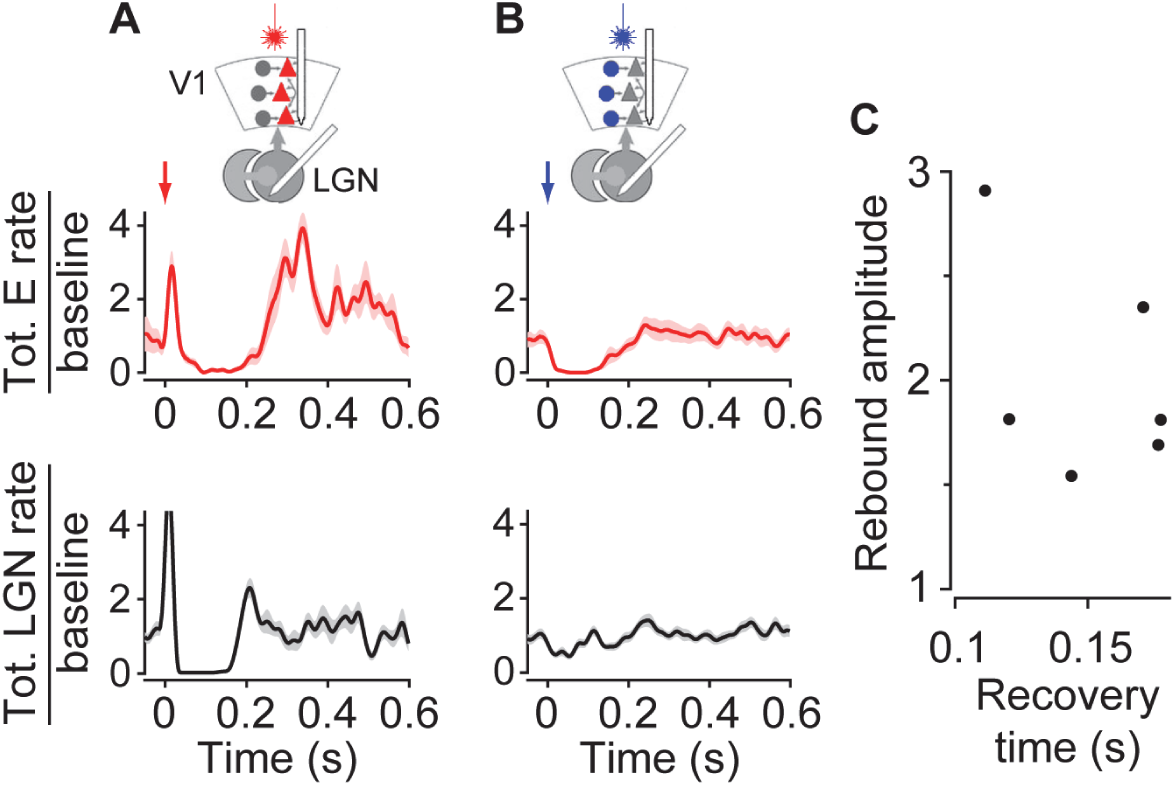
Geniculate and cortical population responses to optogenetic stimulation of V1 excitatory or inhibitory neurons. (A) Peristimulus time histograms of cortical E (red, top) and LGN (black, bottom) population responses (recorded simultaneously) to optogenetic stimulation of cortical E neurons, for an example session. Shaded region: ± s.e.m. across 20 trials. Arrow: optogenetic pulse. (B) As in (A) but for responses to optogenetic stimulation of cortical I neurons. *n* = 35 trials. (C) Recovery time vs. rebound amplitudes of LGN population responses to optogenetic stimulation of cortical E neurons, across sessions. Each point denotes data averaged across all pulse durations in one session. *n* = 6 sessions, 5 mice.

### A corticothalamic model quantitatively predicts evoked cortical and geniculate dynamics

The recordings in LGN suggested a potential explanation for the differences in, and the interactions of, the effects of E and I pulses seen in cortex: whereas an I pulse suppresses cortical firing predominantly via intracortical inhibition, an E pulse additionally drives prolonged suppression followed by a rebound burst in the thalamus. This mechanism can explain the observation that in the paired-pulse experiments, the recovery of activity is locked to the E pulse regardless of the pulse order: a rebound in cortical activity is caused by a rebound in thalamus, which will occur at a fixed time after the E pulse, even if cortical I cells are currently still active. The mechanisms by which an E pulse to the cortex could have this effect on the LGN are well established: corticothalamic neurons provide excitatory input to both thalamic relay cells and the thalamic reticular nucleus (TRN) (Briggs and Usrey, 2008; Sherman and Guillery, 2006); the activation of TRN cells triggers prolonged GABA_A_- and GABA_B_-receptor–mediated inhibition in relay neurons (Huguenard and Prince, 1994; Thomson, 1988). On recovery from this inhibition, voltage-dependent calcium channels in LGN neurons are de-inactivated; subsequent membrane depolarization results in a slow calcium spike that in turn causes burst firing (Halassa et al., 2011; Lu et al., 1992; Scharfman et al., 1990).

We tested whether this hypothesized corticothalamic dynamics could quantitatively explain all our experimental observations by developing a mechanistic corticothalamic (CT) model (**Fig. 7A** and **Supplementary Fig. S7**) that incorporates both thalamic reticular inhibition and burst firing:

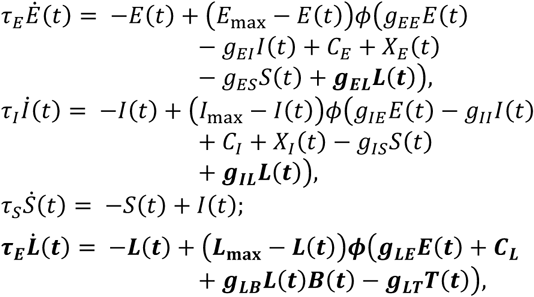

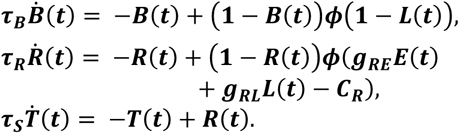

**Fig. 7.**
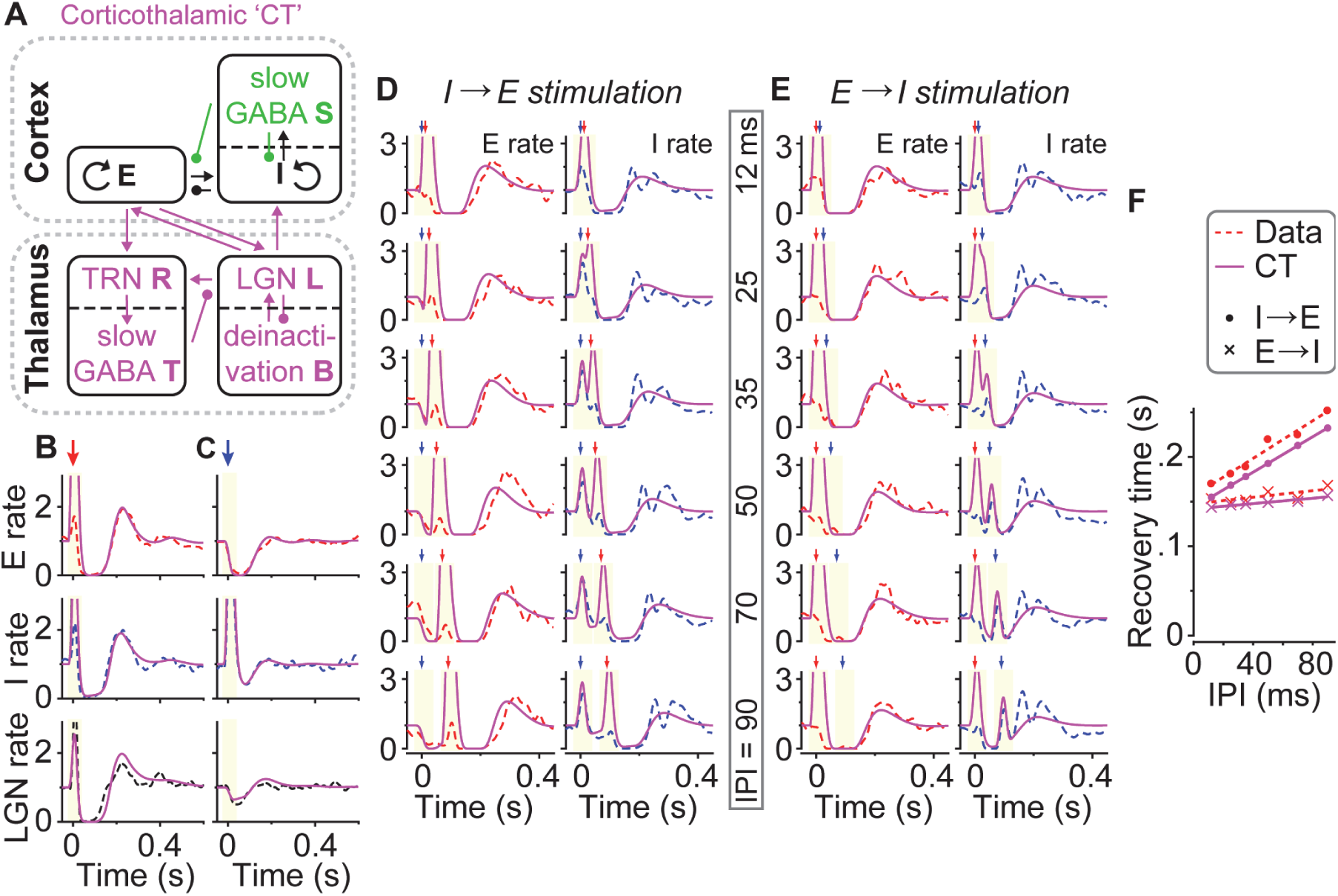
A corticothalamic model quantitatively predicts both cortical and geniculate dynamics. (A) A schematic of the corticothalamic (CT) model. A thalamic component, incorporating thalamic reticular inhibition (*R, T*) and low-threshold relay cell burst firing (*B*), was added to the WCS model. (B,C) Best fit of the CT model to single optogenetic-pulse experimental data. Peristimulus time histograms of experimentally measured E (red dashed, top), I (blue dashed, middle) and LGN (black dashed, bottom) population responses to E (B) and I (C) pulses. Solid magenta lines: CT model predictions. Arrows: optogenetic pulses. Ivory regions: periods excluded in the loss function. Experimental cortical data shown here were averaged across 11 sessions in 11 mice; experimental geniculate data were averaged across 6 sessions in 5 mice; model predictions were averaged across all folds of cross-validation and all sessions. (D,E) Best fit of the CT model to paired-pulse experimental data, for an example session. Red and blue dashed lines: measured E and I population responses, replotted from Fig. 3A,B. Magenta solid lines: CT model predictions. Red and blue arrows: E and I pulses. (F) Recovery time of experimentally measured (red) and CT-predicted (magenta) E population responses to I→E (●) and E→I (×) stimulation as a function of IPI. Red dashed and magenta solid lines are linear fits to data and CT model predictions.

In this set of equations, the terms in **bold** are enhancements over the WCS model. Here, *L*(*t*) and *R*(*t*) are the modeled population rates of LGN and TRN neurons, whose baseline rates prior to stimulation are scaled to 1 and 0, respectively. *T*(*t*) represents slow inhibition resulting from the TRN activity, which, as *S*(*t*), could be understood as representing extra-cellular GABA concentrations or GABA_B_ channel activation. The parameters *g*_*EL*_ and *g*_*IL*_ are the strengths of feedforward connections from the LGN population to the cortical E and I populations. *g*_*LE*_ and *g*_*RE*_ model the strengths of feedback connections from the cortical E population to LGN and TRN. *g*_*RL*_ models the strength of the connection from LGN to TRN; *g*_*LT*_ is the weight of the slow thalamic inhibition on the LGN population; *C*_*L*_ and *C*_*R*_ represent tonic input to LGN and TRN. *B*(*t*) represents the activation state of the voltage-dependent channels responsible for the rebound bursting: as *B*(*t*) grows, it adds a thalamic self-excitatory term *g*_LB_*L*(*t*)*B*(*t*) that models regenerative burst activity, which terminates as *B*(*t*) is driven towards zero when *L*(*t*) ≥ 1. Finally, *g*_LB_ models the propensity of LGN cells toward burst firing.

We fit the parameters of the CT model with the same cross-validated search approach for the purely cortical models. The model fit the experimentally measured dynamics of cortical E and I populations in response to optogenetic stimulation, and we assessed its goodness of fit by the suppression durations and rebound amplitudes.

The CT model provided a good fit for cortical responses to single optogenetic pulses, and key aspects of the modeled responses were statistically indistinguishable from the experimentally measured values (**Fig. 7, 8**). The CT model accurately predicted cortical responses to an E pulse (**Fig. 7B, 8A**; measured and CT-predicted response characteristics across sessions: recovery time, medians = 164 and 165 ms, *p* = 0.32; rebound amplitudes, medians = 2.20 and 2.13 times baseline, *p* = 0.83; *E*_min_, medians = 0 for both, *p* = 0.76; *I*_min_, medians = 0 and 0.02 times baseline, *p* = 0.02; Wilcoxon signed-rank test, 11 sessions in 11 mice). It also accurately predicted cortical responses to an I pulse (**Fig. 7C, 8B**; measured and CT-predicted response characteristics across sessions: recovery time, medians = 123 and 125 ms, *p* = 0.90; rebound amplitudes, medians = 1.36 and 1.24 times baseline, *p* = 0.37; *E*_min_, medians = 0.01 and 0 times baseline, *p* = 0.12; *I*_min_, medians = 0.15 and 0.29 times baseline, *p* = 0.053. Wilcoxon signed-rank test, 11 sessions in 11 mice). Responses to flash visual stimuli were also well fit on a stimulus-by-stimulus basis, using a model modified so that the external visual input went to the LGN instead (Supplemental Information S1.11): the CT model predicted the suppression duration and the minimum E and I population responses post-suppression (measured and CT-predicted response characteristics: recovery time: medians = 146 and 139 ms, *p* = 0.19; *E*_min_, medians = 0.06 and 0.02 times baseline, *p* = 0.30; *I*_min_, medians = 0.09 and 0.05 times baseline, *p* = 0.28; Wilcoxon signed-rank test, 4 conditions per session, 6 sessions in 5 mice). The CT-predicted rebound amplitudes were further improved compared to the WCS model predictions (**Fig. 8C**; CT vs. WCS predictions: *p* = 1×10^−3^, Wilcoxon signed-rank test on absolute difference between experimental data and model predictions, 4 conditions per session, 6 sessions in 5 mice), although still somewhat lower than the experimental data (measured and CT-predicted rebound amplitudes across sessions: medians = 2.39 and 1.97 times baseline, *p* = 7×10^−4^; Wilcoxon signed-rank test, 4 conditions per session, 6 sessions in 5 mice). To further validate the CT model, we tested its ability to predict responses to paired pulses. The CT model captured the observation that recovery post-stimulation was time-locked to the E pulse for both E→I and I→E stimulation (**Fig. 7D–F, 8D,E** and **Supplementary Fig. S6**; CT-predicted slopes for I→E and E→I stimulation across sessions: medians = 0.99 and 0.26, *p* = 4×10^−3^. Data vs. CT-predictions for I→E and E→I stimulation: medians = 1.04 vs. 0.99 and 0.18 vs. 0.26, *p* = 0.20 and 0.30, respectively; Wilcoxon signed-rank test, 9 sessions in 9 mice). The CT model also captured the dynamics of the LGN population; importantly, the model parameters were fit purely to cortical data, suggesting that the model has inferred thalamic dynamics correctly from cortical pulse responses (**Fig. 7B,C**, bottom). We conclude that the CT model provides a concise and accurate account of our experimental observations in both V1 and LGN. This model has more parameters than the purely cortical WC and WCS models, but as the results were cross-validated, tested out-of-sample, and tested on LGN responses while trained only on cortical data, it could not simply gain a numerical advantage by overfitting. The CT model’s superior performance provided a quantitative validation of an intuitive mechanism explaining the difference between the effects of E and I pulses seen in the cortex: prolonged cortical suppression caused by activating cortical E neurons involves a substantial thalamic contribution, whereas the suppression caused by activating cortical I neurons is predominantly due to cortical inhibition.

**Fig. 8.**
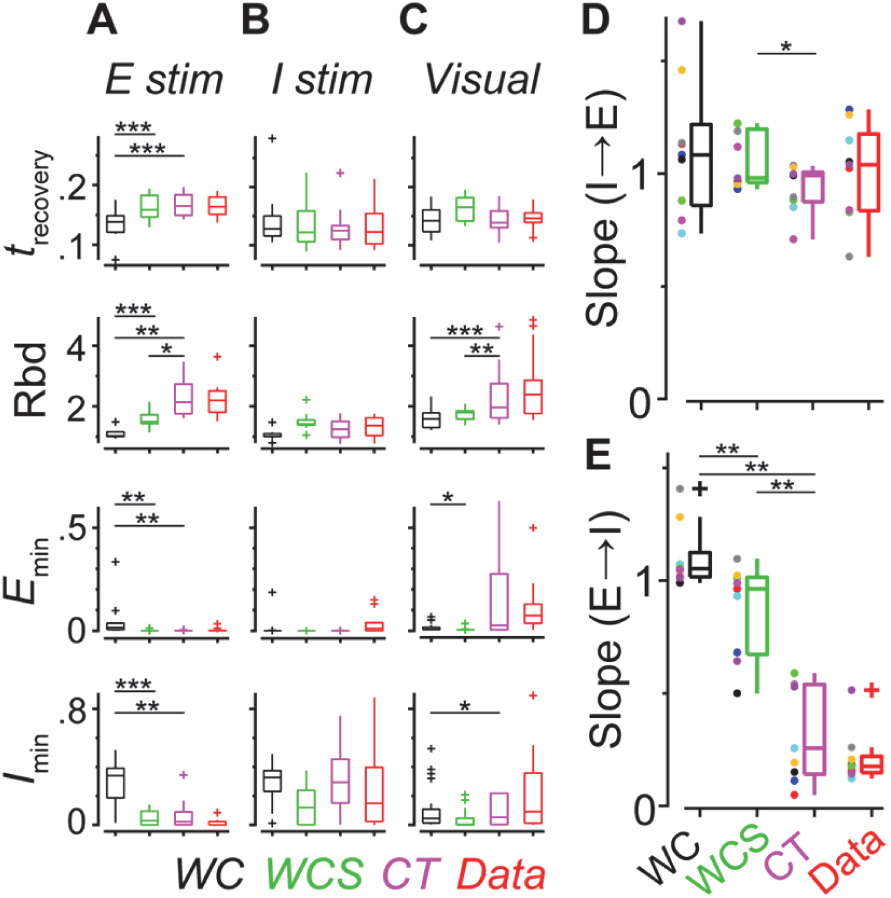
Summary statistics comparing cortical dynamics measured experimentally and predicted by the WC, WCS, and CT models. (A) Box plots summarizing the recovery time (*t*_recovery_), the rebound amplitudes (Rbd), and the minimum E and I population responses (*E*_min_ and *I*_min_) following an E pulse, across sessions. *n* = 11 sessions in 11 mice. Bar, median; box, 25–75% interval; whiskers, 1.5 IQR. Data are in red; WC, WCS, and CT model predictions are in black, green, and magenta. (B) As (A) but for responses to an I pulse. *n* = 11 sessions in 11 mice.(C) As (A) but for responses to flash visual stimuli, across stimulus conditions and sessions. *n* = 4 conditions in a session, 6 sessions in 5 mice. (D) Box plots summarizing the slopes of the linear fits to recovery time as a function of IPI for I→E stimulation. *n* = 9 sessions in 9 mice. Each point marks data from a session. (E) As (D) but for E→I stimulation. Data from the same session share the same color. Wilcoxon signed-rank test on absolute difference between model predictions and experimental data: *, *p* < 0.05, **, *p* < 0.01, ***, *p* < 0.001.

## Discussion

We found that cortical population dynamics in response to optogenetic pulses could be well approximated by a set of mean-field equations, provided that the equations include contributions from the thalamus. The Wilson-Cowan equations could not capture all features of cortical responses to optogenetic pulses, even with an extension that included slow cortical GABAergic inhibition. We could, however, account for the observed dynamics with a corticothalamic model that incorporated prolonged thalamic suppression and rebound bursting activity. The parameters of this model were fit to cortical data alone, but the predicted LGN firing also closely matched the LGN responses measured experimentally.

Our corticothalamic model is highly simplified, taking no account of many well-known processes such as synaptic depression, cellular adaptation, or the existence of multiple excitatory and inhibitory cell types. Understanding the effect of these processes would require more experiments, but it is possible that extending the corticothalamic model to include these features would allow further improvement in its fit to the experimental data. Indeed, while the current model’s predictions are quantitatively accurate, they are not perfect. For example, after the suppression and rebound evoked by an E pulse, the network activity decays to a value slightly below its baseline level (the same is not true of an I pulse). This experimental finding is not captured by any of our current models. A slowly recovering synaptic depression might be able to model this behavior by shifting the fixed point of the excitatory-inhibitory network. Another discrepancy is the response to paired EE pulses (**Supplementary Fig. S6**), which was modeled less accurately than any other paired-pulse combination: according to the models, the recovery time is determined solely by the timing of the second pulse, but the experimental data also showed a small effect from the timing of the first pulse. Finally, while there are many classes of cortical inhibitory cells, our models only contained one of them – the parvalbumin-expressing interneurons – whose activity was also the only class we were able to measure. Other interneuron classes, such as neurogliaform cells, can produce long-lasting hyperpolarizing currents via slow non-synaptic inhibition (Overstreet-Wadiche and McBain, 2015; Simon et al., 2005). Nevertheless, we found that simply adding a term for slow cortical inhibition to the Wilson-Cowan equations was insufficient to predict quantitatively all our experimental observations. A technical limitation of our recording method is that it severely underestimated the firing rates within ∼20 ms immediately following an E pulse, owing to spike overlaps and fast local field potential changes. Consequently, we could not use the magnitude of the initial response to constrain our models. New technical developments, such as accurate voltage indicators (Abdelfattah et al., 2019; Adam et al., 2019; Fan et al., 2019), might allow accurate measurements of the responses of multiple cell types even during periods of high synchrony, providing additional constraints on the models.

Our results indicate that the type of suppression evoked by an I pulse is fundamentally distinct from the type of suppression evoked by an E pulse. This was suggested by observations such as the order-dependent responses to paired pulses, which could not be accurately described by purely cortical models. Even stronger evidence was provided by recordings in the LGN, which showed strong thalamic suppression and rebound following an E (but not I) pulse to the cortex, and by the closer match of predictions from a corticothalamic model to the data. The models suggest that following an I pulse to the cortex, both cortical E and I populations are suppressed because inhibition of E neurons removes drive from both neuronal classes. This is essentially the mechanism of an inhibitory-stabilized network (Li et al., 2019; Mahrach et al., 2020; Tsodyks et al., 1997); indeed, our cortical models were of this type. Interestingly, the parameters of the corticothalamic model did not always show strong enough cortical excitation to alone qualify as ISN, yet when recurrent activity via thalamus was included, we again observed ISN-like behavior. This suggests that cortical suppression following an I pulse to the cortex might also have a thalamic contribution: because an I pulse suppresses cortical E firing, it indirectly suppresses thalamic firing owing to lower feedback input, which in turn lowers the amount of feedforward input received by cortical cells. Indeed, thalamic activity is slightly suppressed following an I pulse to the cortex (although the magnitude of the suppression is significantly less than following an E pulse) (**Fig. 6B**). It is thus possible that the strong recurrent excitation required for the ISN-like behavior arises not just from cortical synapses, but also via loops through thalamus and perhaps also other brain regions. In contrast, the suppression in response to an E pulse to the cortex, with its longer duration and much stronger rebound on recovery, appears much more like a single cycle of the well characterized classical thalamic oscillation that occurs also during sleep spindles (Destexhe and Sejnowski, 2003; McCormick and Bal, 1997), for example.

Prolonged suppression of thalamocortical activity did not only result from direct optogenetic stimulation of cortical neurons but could also occur in response to visual stimuli. This result is consistent with previous findings from the visual system of mice and humans (Funayama et al., 2016; Minamisawa et al., 2017), as well as the auditory system of mice and cats (Guo et al., 2017a; Mariotti et al., 1989), where sensory stimuli also evoked prolonged thalamic suppression. Prolonged thalamic suppression, leading in turn to prolonged cortical suppression followed by rebound activity, is thus a common response to strong activation of cortex or thalamus. The computational role of these prolonged suppressions is not clear. One clue might come from that they are stronger during sleep (Mariotti et al., 1989), forming a potential mechanism for the large “K-complex” wave seen in human scalp recordings (Cash et al., 2009). These prolonged inhibitions might therefore serve to prevent strong sensory stimuli from “overloading” the cortex in less attentive states. Alternatively, or additionally, it is possible that the rebound burst following a spindle provides a carefully timed window for enhanced sensory processing (Funayama et al., 2016; Guo et al., 2017a; Minamisawa et al., 2017). We propose that a similar mechanism of prolonged suppression evoked by externally induced bursts in cortical pyramidal cells underlies the ability of cortical electrical stimulation (Chung and Ferster, 1998) to suppress cortical activity, and speculate that a similar phenomenon might explain how pulsatile transcranial magnetic stimulation transiently suppresses cortical activity. In summary, we found that cortical dynamics can be well approximated by a simple mean-field model, but only if this model included contributions from the thalamus. The equations we have derived provide a good fit to our experimental data, but they can be further constrained and refined by additional experiments measuring activities of multiple interneuron types and using recording techniques capable of accurately measuring the initial response to brief, strong optogenetic stimulation. Such models would constitute a first step towards a “statistical mechanics of the cortex”.

## Experimental Procedures

All experiments were conducted under personal and project licenses issued by the Home Office, in accordance with the UK Animals (Scientific Procedures) Act 1986. Experimental methods are detailed in Supplemental Information S1.

## Supplemental Information

Supplemental Information includes Supplemental Experimental Procedures, seven figures, and four tables. It can be found with this article online at X.

## Acknowledgments

This work was supported by the Wellcome Trust (grant 205093), the Simons Collaboration on the Global Brain (grant 325512), and the European Research Council (grant 694401). M.C. holds the GlaxoSmithKline/Fight for Sight Chair in Visual Neuroscience. M.O. is funded by the Academy of Medical Sciences, the Wellcome Trust (grant SBF002/1045), and BBSRC (grant BB/P020607/1).

## Supplemental Information

**Fig. S1.**
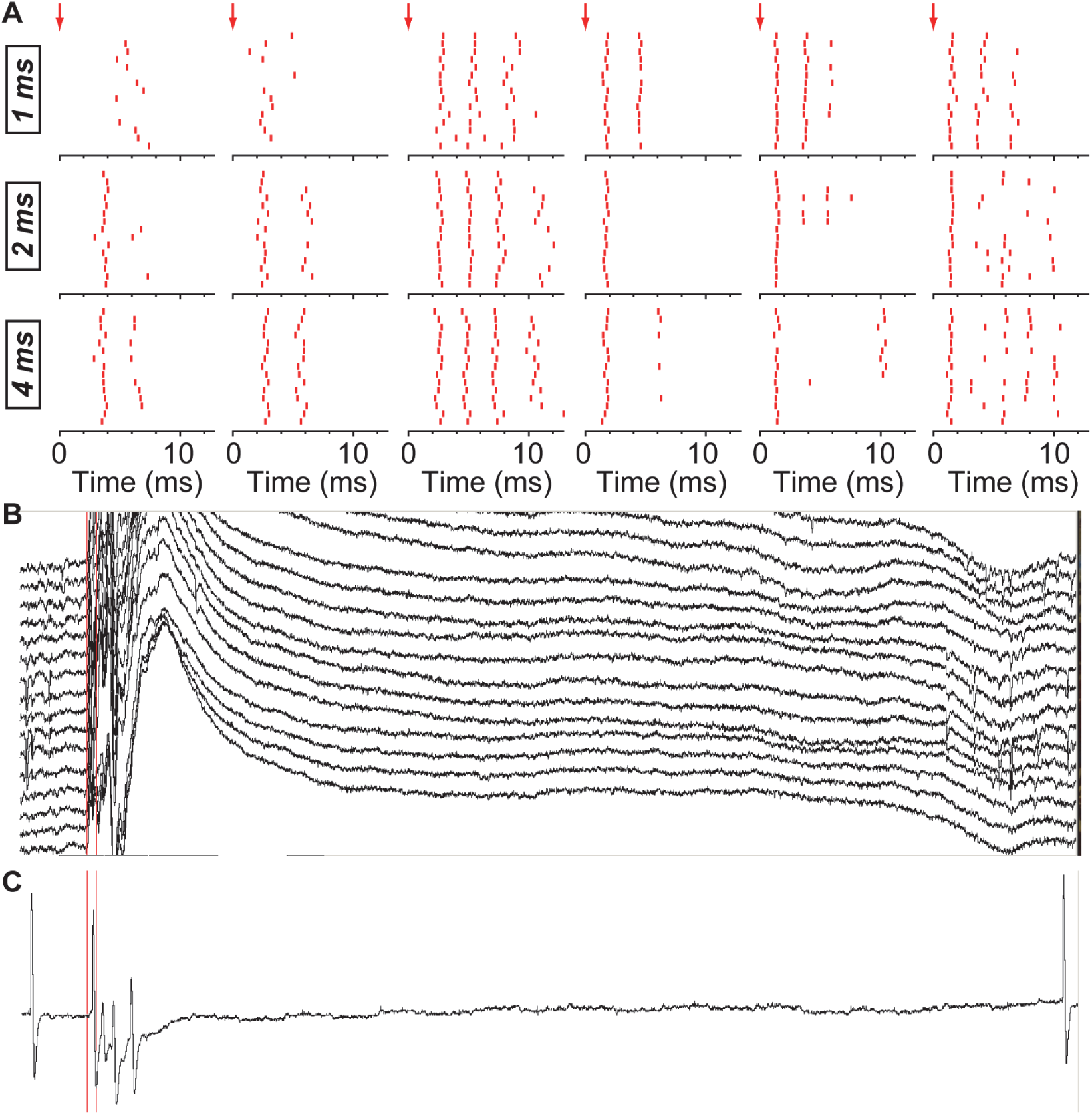
Juxtacellular and extracellular recordings of cortical responses to optogenetic stimulation of V1 excitatory neurons. (A) Raster plots of initial responses of 6 example cortical E neurons to 15 repeats of 1-, 2-, and 4-ms E pulses. The onset of the pulse is marked by a red arrow. Trials of different pulse durations were randomly interleaved but are displayed here as separate rasters. Note that spiking could occur as early as 1 ms after pulse onset. (B) Raw electrophysiological data traces (duration = 225 ms) recorded with a Buzsaki32 probe (only data from 16 recording sites are shown here), following a 2-ms E pulse (red bar). Note that overlapping spikes and fast local field potential oscillations preclude spike sorting for ∼20 ms after the pulse onset. (C) Raw electrophysiological trace (duration = 225 ms) of a juxtacellular electrode from a separate recording. The trace shows one spike preceding the E pulse and four spikes in response to the E pulse, with the first of the four happening fast enough to overlap with the pulse.

**Fig. S2.**
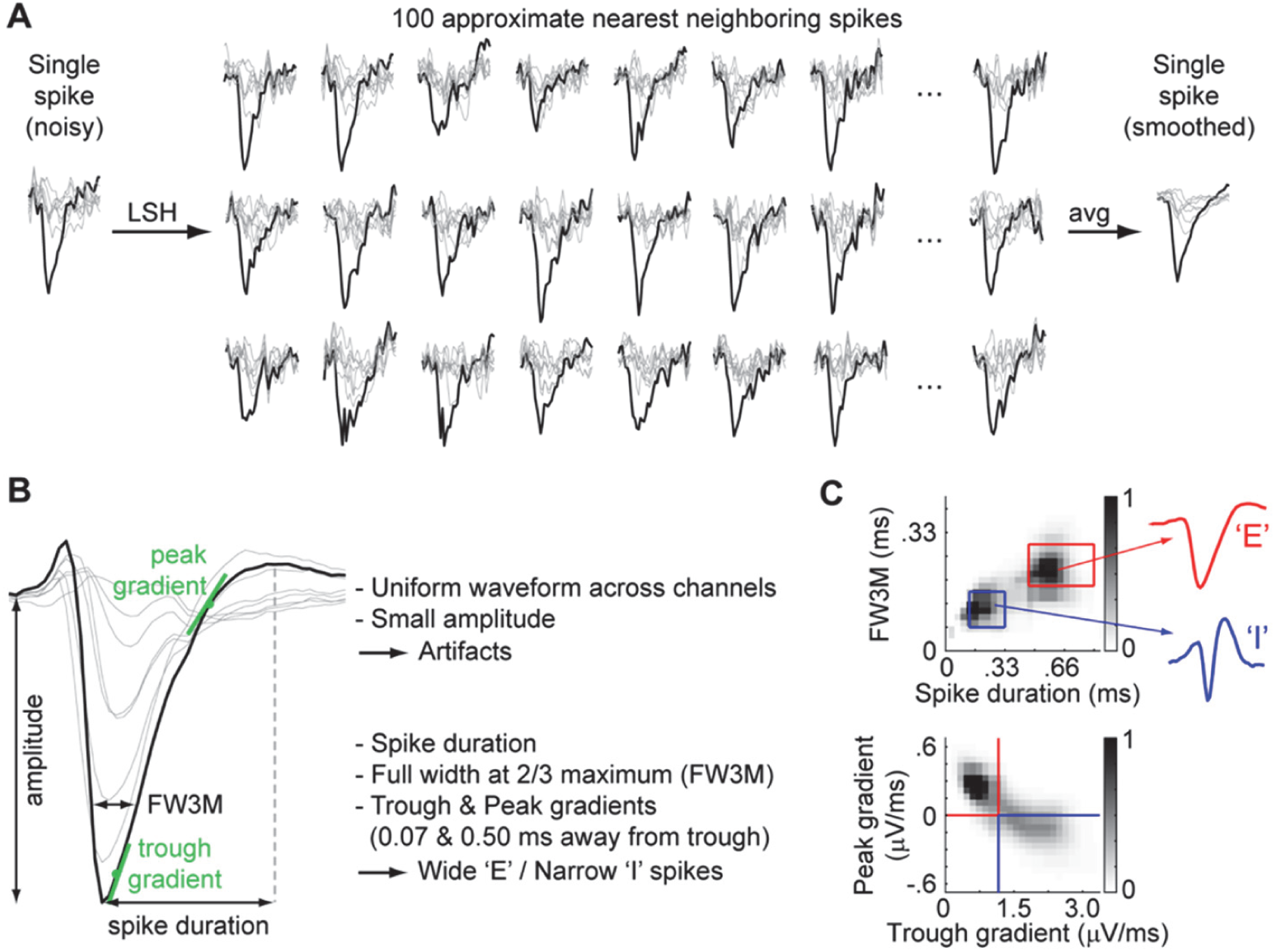
A “clusterless” method to separate wide and narrow spikes. (A) To allow accurate measurement of spike-shape characteristics, raw spikes are first denoised by averaging over spikes in similar regions of feature space. To do this, for each target spike, we used locality-sensitive hashing (LSH) to find 100 spikes of similar waveform on all recording sites ‘channels’ of the same shank; we then averaged the waveforms of these 100 spikes to obtain a smoothed waveform for the target spike. The example raw spike (left) is taken from a recording with a Buzsaki32 probe. The black trace is from the channel with maximum spike amplitude; the gray traces are from the other seven channels of the same shank. The middle panels show few example ‘neighboring’ spikes, and the right panel shows the denoised spike. (B) Waveform characteristics used to separate wide (putative excitatory) and narrow (putative fast-spiking inhibitory) spikes: full-width at 2/3 maximum (FW3M), trough-to-peak duration; early and late gradients of the upstroke (measured at 0.07 and 0.50 ms from the trough time). The spike amplitude and the cross-channel waveform variability were also used for quality control: spikes of low amplitude or of similar waveforms across channels were excluded from further analysis. (C) Top: density plot of FW3M vs. spike duration for all spikes in an example recording session. Bottom: density plot of peak gradient vs. trough gradient. The boundaries to classify putative excitatory and fast-spiking inhibitory spikes were marked in red and blue.

**Fig. S3.**
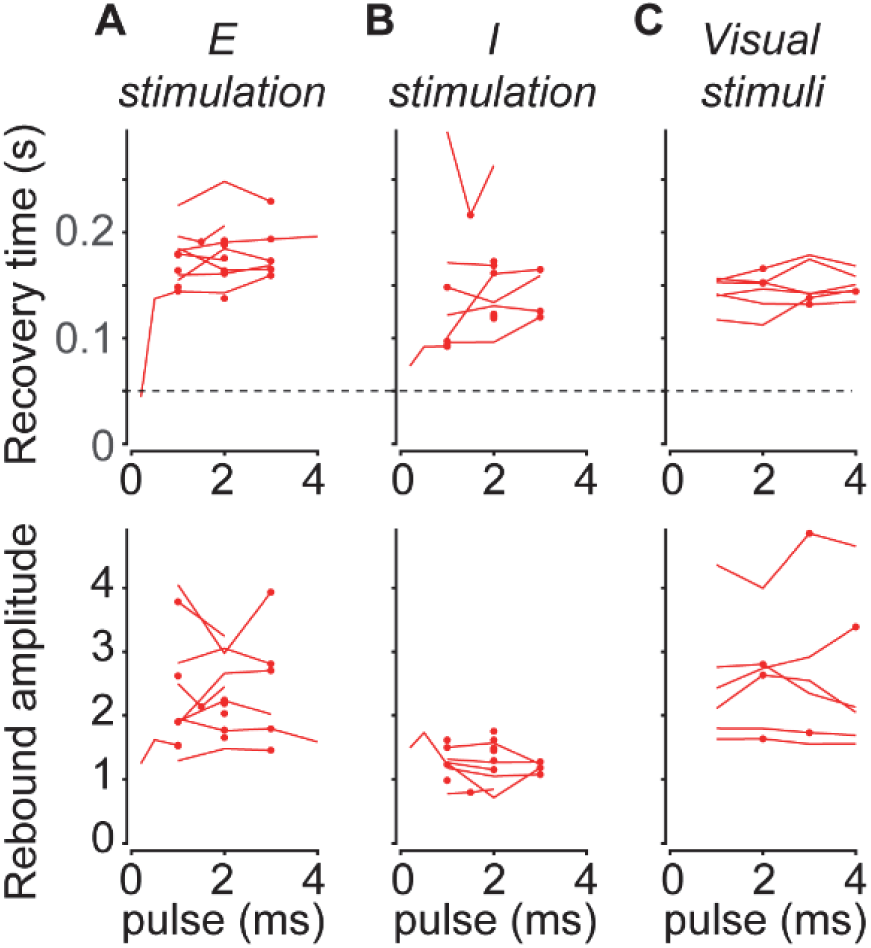
Dependence of impulse-response cortical dynamics on durations of E and I pulses as well as flash visual stimuli. (A) Recovery time (top) and rebound amplitudes (bottom) of E population responses to single E pulses as a function of pulse duration. As before, recovery time is the time when the E population response recovers to 50% of its baseline activity; rebound amplitude is the peak E population response post-suppression. Lines connect data from the same recording session (9 out of 17 sessions had multiple pulse durations); dots marked the data plotted in Fig. 2A. The dashed line marked the threshold of 50 ms; for Fig. 2D, we did not include data in which the applied optogenetic pulse was not strong enough to evoke a suppression > 50 ms (only one dataset, with pulse length = 0.2 ms). (B) As (A) but for responses to single I pulses. 7 out of 14 sessions had multiple pulse durations. (C) As (A) but for responses to a brief light flash to the contralateral eye. *n* = 6 sessions, 5 mice. All sessions had multiple flash durations.

**Fig. S4.**
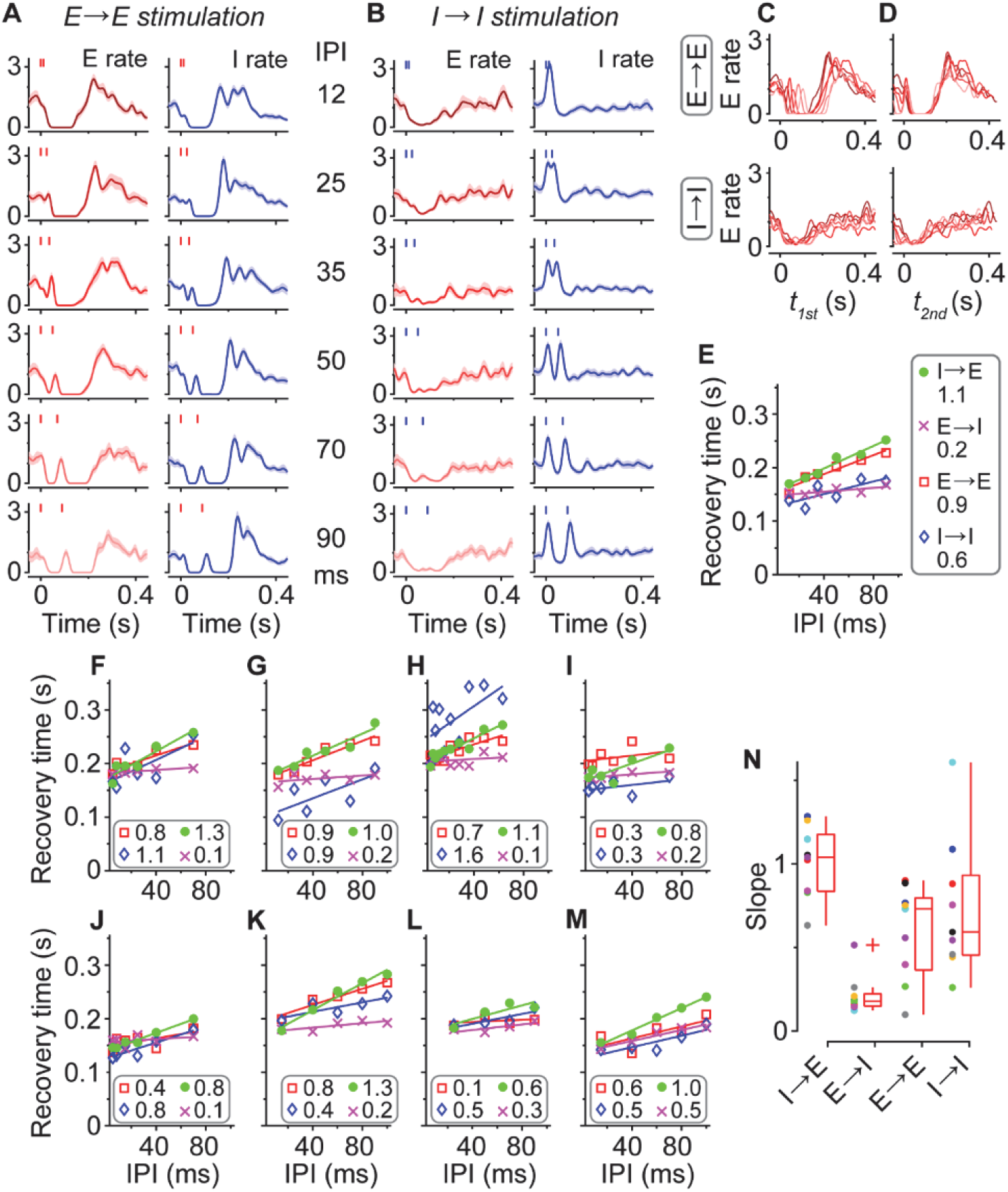
Population responses to paired EE and II optogenetic pulses. (A) Peristimulus time histograms of cortical E and I population responses to paired E pulses (E→E stimulation) at various interpulse intervals (IPIs), for an example session. Red trace (left column): mean E population rate; Blue trace (right column): mean I population rate. Each row shows the mean response across trials with a fixed IPI, normalized so baseline rate is 1. Short red and blue lines: E and I pulses. Shaded area: s.e.m. across trials in a session. (B) As in (A) but for responses to paired I pulses (I→I stimulation). (C) E population responses to E→E (top) and I→I (bottom) stimulation for the same session, overlaid and re-aligned to the first pulse. Colors represent the IPIs, matching rows in (A). (D) As in (C) but aligned to the second pulse. (E) Recovery time of E population responses to I→E (●), E→I (×), E→E (□), and I→I (◁) stimulation as a function of IPI, for this example session. Solid lines, linear fits; the slopes of the fits are noted in the legend. (F–M) As in (E) but for the other 8 recording sessions. (N) Box plots summarizing the slopes of the linear fits to recovery time vs. IPI up to 100 ms across 9 sessions in 9 mice. Bar, median; box, 25–75% interval; whiskers, 1.5 IQR. Each dot marks data from a session; data from the same session share the same color.

**Fig. S5.**
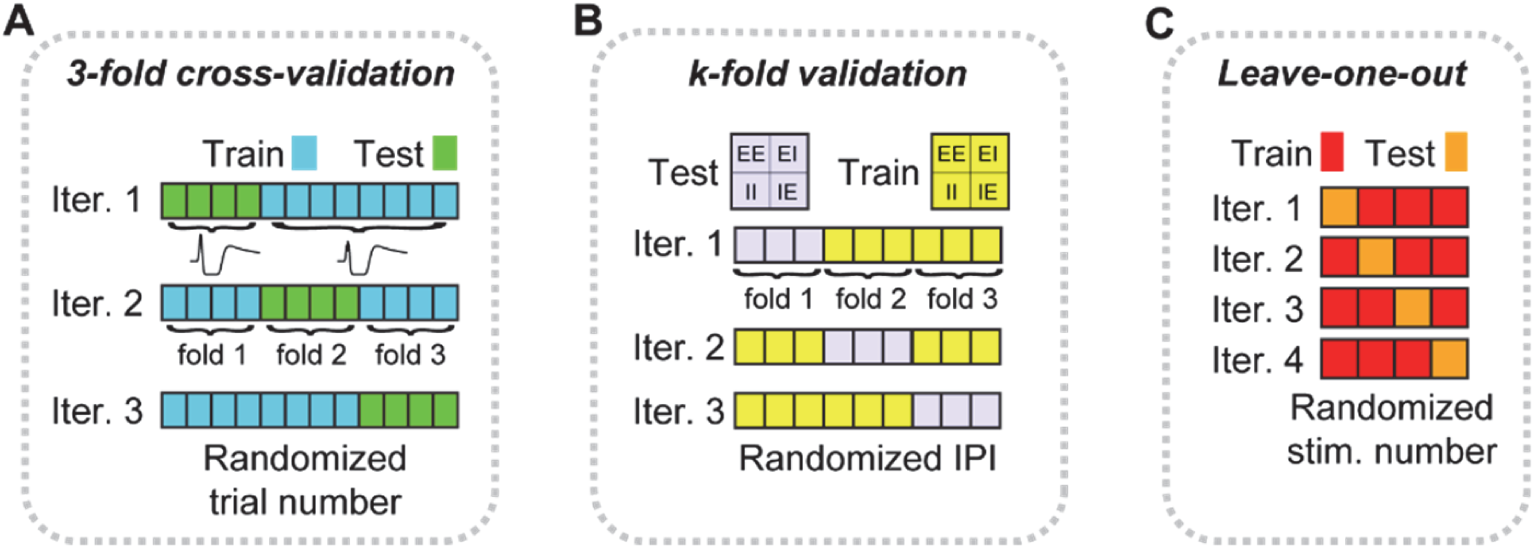
Validation methods. (A) A schematic of the 3-fold cross-validation method used to evaluate models for responses to single E and I pulses. First, we randomly partitioned the trials into three folds: for a recording session with *k* trials of E pulses and *k* trials of I pulses, each fold had *k/3* randomly chosen trials of E pulses and *k/3* randomly chosen trials of I pulses. Each fold took turn as the test set. We trained the model on peristimulus time histograms (PSTHs) averaged across trials of the other two training-set folds (blue) and assessed its performance on PSTHs averaged across trials of the test fold (green). (B) A schematic of the out-of-sample validation method used to assess responses to paired pulses. We randomly divided the *N* interpulse intervals (IPIs) into *k* folds (*k* = 2 or 3, depending on the total number of IPIs; see Supplementary Table S3) so that each fold contains *N/k* IPIs. For each of these folds, we trained the model on the other *k-1* folds (yellow, ‘train’) and assessed the model’s performance on the remaining fold (purple, ‘test’). Note that for every IPI, there are four paired-pulse combinations: paired E pulses (‘EE’), paired I pulses (‘II’), first E then I pulse (‘EI’), and first I then E pulse (‘IE’). Thus, for every iteration, the model was trained on 4(*k*-1)*N*/*k* conditions, and its performance was assessed on the remaining 4*N*/*k* conditions. (C) A schematic of the out-of-sample method adopted to fit models to responses to flash visual stimuli. There are four stimulus conditions (1-, 2-, 3-, and 4-ms flash durations) in each recording. For each condition, we trained the model on the trial-averaged PSTHs of the other three conditions (red, ‘train’) and assessed its performance in predicting the trial-averaged PSTH for the remaining condition (orange, ‘test’). These steps were repeated for each of the four conditions as the test set.

**Fig. S6.**
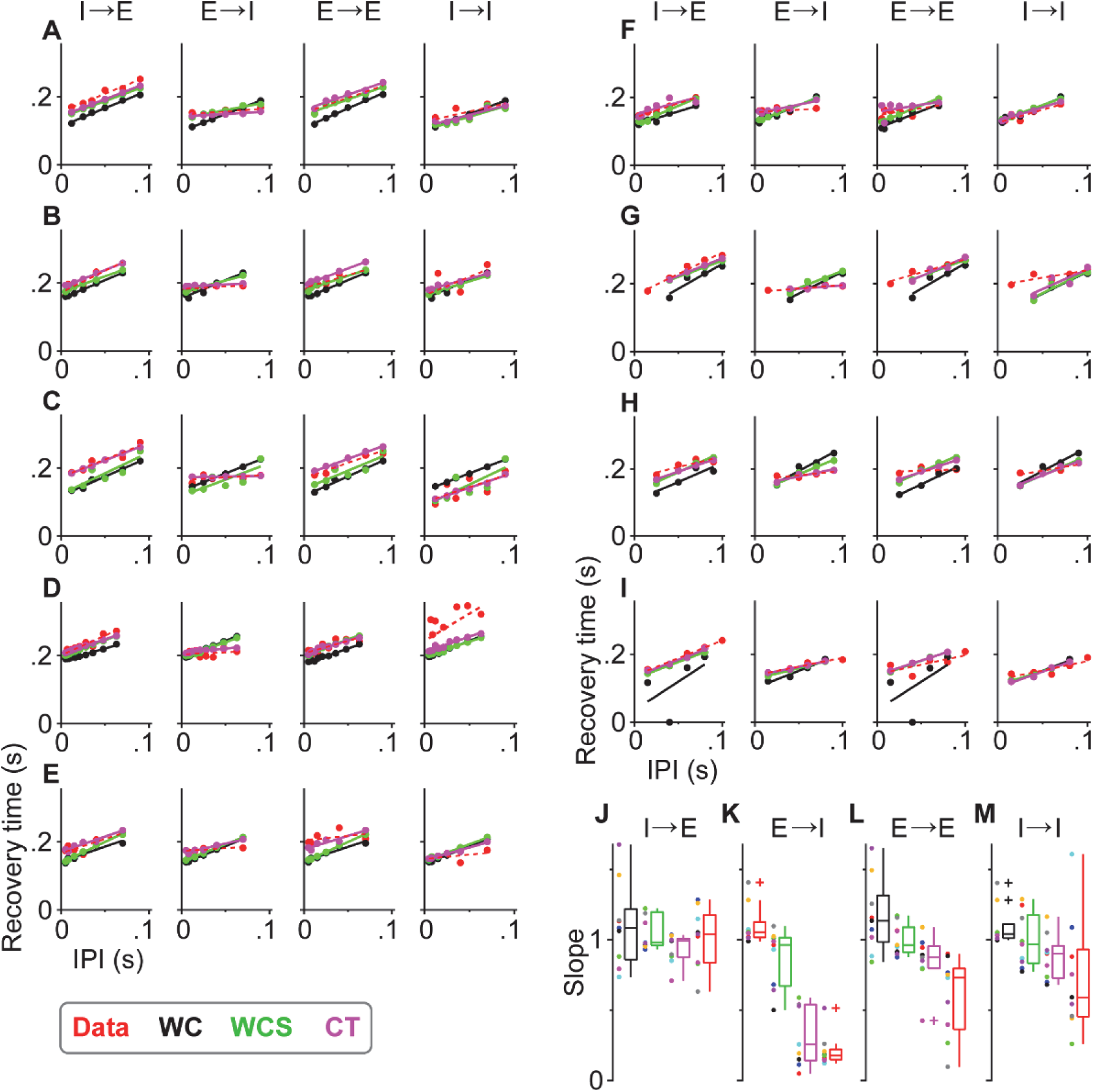
Comparisons between measured and modeled cortical responses for all sessions and paired-pulse combinations. (A) Recovery time of measured and modeled E population responses to all four paired-pulse combinations as a function of IPI, for the example session in Fig. 3A,B. Red points and dashed line represents experimental data; black: WC model, green: WCS model, magenta: CT model. Red, black, green, and magenta solid lines are linear fits to data and WC, WCS, and CT model predictions. (B–I) As in (A) but for the other 8 recording sessions. (J) Box plots summarizing the slopes of the linear fits to recovery time as a function of IPI for I→E stimulation. *n* = 9 sessions in 9 mice. Bar, median; box, 25–75% interval; whiskers, 1.5 IQR. Data are in red; WC, WCS, and CT model predictions are in black, green, and magenta. Each point marks data from a session; data from the same session share the same color. (K–M) As in (J) but for E→I (K), E→E (L), and I→I (M) stimulation. Note the large variability between experiments for E→E and I→I stimulation.

**Fig. S7.**
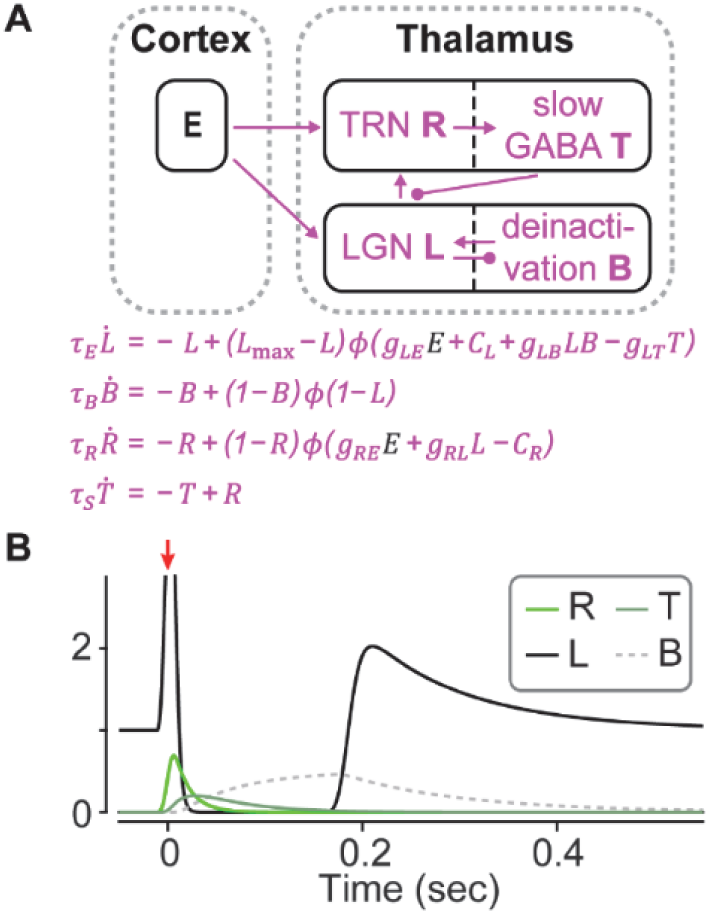
A model of prolonged thalamic inhibition and rebound bursting firing. (A) Cortical E neurons provide feedback excitatory inputs to both LGN (*L*) and TRN (*R*), whose activation drives slow thalamic GABAergic inhibition (*T*). The de-inactivation parameter *B* models the activation state of the voltage-dependent channels responsible for rebound bursting. (B) Dynamics of the four thalamic variables following an E pulse to the cortex (onset marked by a red arrow). Note that the value of *B* starts to increase as soon as *L* dips below 1, yet due to its appearance in a product term *LB* it does not cause rebound bursting until *L* begins to increase from zero.

**Table S1.** Mouse strains, the opsins they expressed (if any), and the corresponding laser wavelength to activate the opsin.

| Strain + AAV vector | Pyramidal (E) |  | Parvalbumin-expressing (I) |  |
| --- | --- | --- | --- | --- |
|  | Opsin | Wavelength | Opsin | Wavelength |
| C57BL/6J | - | - | - | - |
| Thy18 | ChR2 | 445 nm | - | - |
| PV<sup>cre</sup>;Ai32 | - | - | ChR2 | 445 nm |
| PV<sup>cre</sup>;Thy18 + C1V1<sub>T/T</sub> | ChR2 | 445 nm | C1V1<sub>T/T</sub> | 561 nm |

**Table S2.** Experiment details, including mouse ID, genotype, amount of virus injected (if any), sex, the depth of each neural probe’s tip below the brain surface (inserted straight down), and modes of stimulation (together with number of trials) for each animal. Buzsaki32 silicon probes were used unless otherwise stated. ^a^ No virus expression; ^b^ simultaneous recordings in the V1 and LGN.

| Mouse ID | Genotype | Virus (nl) | Sex | V1 depth (μm) | LGN depth (μm) | Single E pulse |  | Single I pulse |  | Paired pulses | Light flash |
| --- | --- | --- | --- | --- | --- | --- | --- | --- | --- | --- | --- |
|  |  |  |  |  |  | Number of trials per stimulus condition |  |  |  |  |  |
| M150605A | PV<sup>cre</sup>;Thy18 | 506 | F | 591 | - | 31 |  | 31 |  | 22 | - |
| M150909C | PV<sup>cre</sup>;Thy18 | 437 | F | 615 | - | 15 |  | 15 |  | 15 | - |
| M150823B | PV<sup>cre</sup>;Thy18 | 414 | F | 605 | - | 22 |  | 22 |  | 22 | - |
| M150303B | PV<sup>cre</sup>;Thy18 | 483 | F | 535 | - | 15 |  | 15 |  | 10 | - |
| M150609A | PV<sup>cre</sup>;Thy18 | 414 | F | 550 | - | 68 |  | 68 |  | 30 | - |
| M150609B | PV<sup>cre</sup>;Thy18 | 414 | F | 605 | - | 61 |  | 61 |  | 28 | - |
| M161122 | PV<sup>cre</sup>;Thy18 | 425 | F | 600 | - | 20 |  | 20 |  | 20 | - |
| M151020A | PV<sup>cre</sup>;Thy18 | 437 | M | 700 | - | 30 |  | 30 |  | 20 | - |
| M160817B | PV<sup>cre</sup>;Thy18 | 414 | M | 845 (Edge32) | 2775 | 15 (V1) | 20 (LGN) | 15 (V1) | 20 (LGN) | 20 | - |
| M141020A | PV<sup>cre</sup>;Thy18 | 495 | F | 550 | - | 40 |  | 40 |  | - | - |
| M160817A | PV<sup>cre</sup>;Thy18 | 414 | M | 774 | - | 25 |  | 20 |  | - | - |
| M150909A | PV<sup>cre</sup>;Thy18 | 437<sup>a</sup> | F | 600 | - | 35 |  | - |  | - | - |
| M150909B | PV<sup>cre</sup>;Thy18 | 437<sup>a</sup> | F | 610 | - | 20 |  | - |  | - | - |
| M150309A | PV<sup>cre</sup>;Thy18 | 451<sup>a</sup> | F | 620 | - | 18 |  | - |  | - | - |
| M150224 | Thy18 | - | F | 585 | - | 55 |  | - |  | - | - |
| M160802<sup>b</sup> | Thy18 | - | F | 750 | 2725 | 35 |  | - |  | - | - |
| M160908<sup>b</sup> | Thy18 | - | F | 665 | 2675 | 20 |  | - |  | - | - |
| M160804 | Thy18 | - | F | - | 2565 | 15 |  | - |  | - | - |
| M151120B | Thy18 | - | F | - | 2950, 2585 | 20, 15 |  | - |  | - | - |
| M160906<sup>b</sup> | PV<sup>cre</sup>;Ai32 | - | F | 720, 600 | 2655, 2575 | - |  | 35 |  | - | - |
| M160921<sup>b</sup> | PV<sup>cre</sup>;Ai32 | - | F | 619 | 2445 | - |  | 20 |  | - | - |
| M160226 | PV<sup>cre</sup>;Ai32 | - | M | - | 2715 | - |  | 10 |  | - | - |
| M151105 | PV<sup>cre</sup>;Ai32 | - | F | - | 3000 (Edge32) | - |  | 12 |  | - | - |
| M160210A | C57BL/6J | - | F | 600 | 2760, 2845 | - |  | - |  | - | 16 |
| M160210B | C57BL/6J | - | F | 675 | 2450, 2320 | - |  | - |  | - | 24 |
| M181130B | C57BL/6J | - | F | 700 | - | - |  | - |  | - | 30 |
| M181130C | C57BL/6J | - | F | 750 | - | - |  | - |  | - | 20 |
| M181204B | C57BL/6J | - | F | 600, 650 | - | - |  | - |  | - | 30 |
| M170308 | Thy18 | - | M | Juxta | - | Y |  | - |  | - | - |
| M170522 | Thy18 | - | M | Juxta | - | Y |  | - |  | - | - |
| M170602 | Thy18 | - | M | Juxta | - | Y |  | - |  | - | - |
| M170614 | Thy18 | - | F | Juxta | - | Y |  | - |  | - | - |

**Table S3.** The interpulse intervals (IPIs) used in each recording session (labelled by the mouse ID) with paired-pulse stimuli. Also tabulated is the number of folds used in the *k*-fold out-of-sample validation method (see Supplemental Information S1.10 and Supplementary Figure S5) for fitting models to the pulse-pair data in each session. IPIs in brackets were not included in model fitting.

| Mouse ID | Interpulse interval (ms) | *k*-fold |
| --- | --- | --- |
| M150605A | 5, 8, 15, 25, 40, 70, 120, 205, 350, (590, 800) | 3 |
| M150909C | 12, 25, 35, 50, 70, 90, 120, 200 | 2 |
| M150823B | 12, 25, 35, 50, 70, 90, 120, 200 | 2 |
| M150303B | 5, 7, 9, 12, 16, 21, 28, 36, 48, 63, 110, 145, 191, 251, 331, 437, (575, 759) | 2 |
| M150609A | 5, 8, 15, 25, 40, 70, 120, 205, 350, (590, 800) | 3 |
| M150609B | 5, 8, 15, 25, 40, 70, 120, 205, 350, (590, 800) | 3 |
| M161122 | (15), 40, 60, 80, 100, 120, 205 | 3 |
| M151020A | 25, 50, 70, 90, 120, 200 | 3 |
| M160817B | 15, 40, 60, 80, 100, 120, (205) | 3 |

**Table S4.**
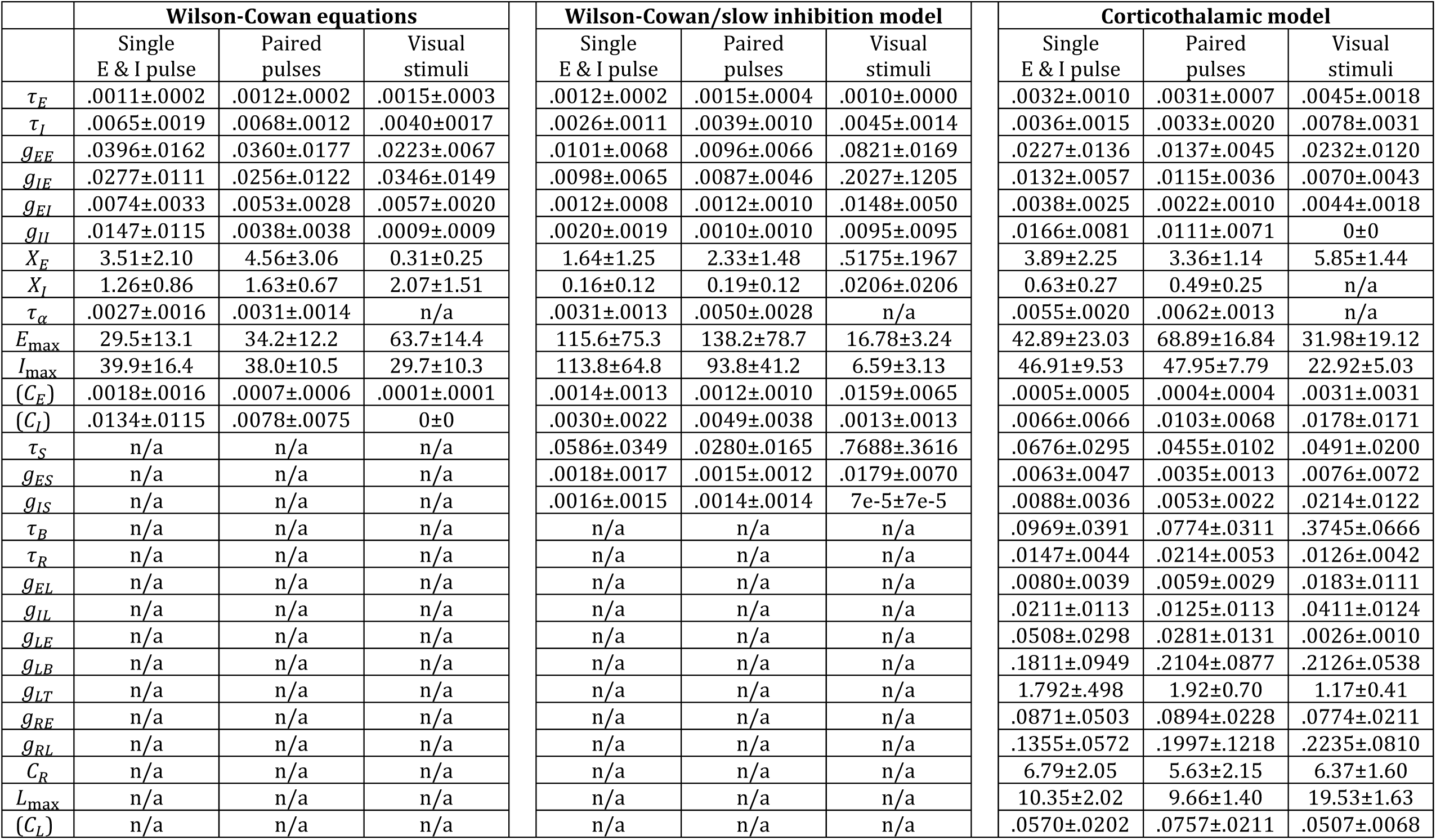
Median ± MAD of the model parameters fitted using cross-validation, adopted to the three experimental paradigms we used: single optogenetic pulse, paired pulses, and flash visual stimuli. There are 33 sets of parameters for single pulses (3 folds x 11 recording sessions), 24 sets for paired pulses (various folds, 9 sessions, see Table S3), and 24 sets for flash visual stimuli (4 stimulus conditions x 6 sessions). Parameters in brackets were not fitted but inferred analytically.

## S1. Supplemental Experimental Procedures

All experiments were conducted according to the UK Animals Scientific Procedures Act (1986), under personal and project licenses released by the Home Office following appropriate ethics review.

### S1.1 Mouse strains

We recorded from adult mice (aged 10–28 weeks at the time of the first recording). Double transgenic mice were generated by crossing B6;129P2-*Pvalb*^*tm1(cre)Arbr*^/J (PV^cre^, #008069; Hippenmeyer et al. (2005)) with either B6;129S-*Gt(ROSA)26Sor*^*tm32(CAG-COP4*H134R/EYFP)Hze*^/J (Ai32, #012569; Madisen et al. (2012)) or B6.Cg-Tg(Thy1-COP4/EYFP)18Gfng/J mice (Thy1-ChR2-YFP founder line18 ‘Thy18’, #007612; Arenkiel et al. (2007)). PV^cre^ mice express Cre recombinase in parvalbumin-expressing neurons (fast-spiking I cells). Ai32 mice express ChR2 following exposure to Cre recombinase. Thy18 mice express ChR2 in layer V cortical E neurons under the control of the Thy1 promotor. All strains were obtained from the Jackson Laboratory, USA.

We recorded from five C57BL/6J wild-type mice, ten Thy18 mice, four PV^cre^;Ai32 mice, and fourteen PV^cre^;Thy18 mice injected with C1V1_T/T_ conditional virus (**Tables S1** and **S2**).

### S1.2 Head-plate implant

Using aseptic techniques, each mouse was implanted with a custom-built head-plate with a recording chamber under isoflurane anesthesia. During the surgery, eyes were protected with ophthalmic gel (Viscotears Liquid Gel, Alcan), and the core body temperature was maintained at around 37°C. After surgery, the animal was provided with rimadyl in drinking water for 3 days.

### S1.3 Virus injections

Stereotaxic injections were carried out in PV^cre^;Thy18 mice. A total volume of 414–506 nl of virus solution (AAV9.Ef1a.DIO.C1V1(E122T/E162T).TS.eYFP.WPRE.hGH, UPenn Vector Core; titer: 5.86e11 GC/ml) was unilaterally injected into the left V1 (approximately 3.2 mm posterior and 2.6 mm lateral from the bregma; 0.5 mm below the exposed dura surface). Injections were made using a glass micropipette (Drummond Scientific, USA) pulled with a micropipette puller (Sutter Instrument, USA) to form a long narrow tip (diameter 25–30 µm). 2.3 nl of the virus solution was injected every 6 seconds; the micropipette was withdrawn 10 minutes after completing the injection. We allowed at least 4 weeks for the virus to express. Note that due to transient developmental expression of parvalbumin, ectopic Cre expression may be found in a small subset of excitatory neurons (Moore and Wehr, 2013; Tanahira et al., 2009).

### S1.4 Juxtacellular and extracellular recordings in awake mice

After three head-restraint acclimatization sessions, the animal was briefly anesthetized with 1.5–2% isoflurane, and a small craniectomy was made over the left V1 (approximately 3.7 mm posterior and 3.0 mm lateral from the bregma) and/or LGN (approximately 2.3 mm posterior and 2.1 mm lateral from the bregma). When necessary, the dura was resected with a 30G needle. The brain was covered with cortex buffer and sealed using Kwik-Cast (World Precision Instruments, USA); the animal was allowed to recover for at least 2 hours before recording. If additional recordings were carried out on subsequent days, the brain was again protected with cortex buffer and Kwik-Cast at the end of each recording session. During a session that typically lasted for 3 hours, the animal stood in a custom-built tube and was judged to be quietly awake by video monitoring (open eyes, postural adjustments, as well as sporadic whisking and grooming).

Juxtacellular recordings of 50 neurons were performed in four Thy18 mice (**Table S2**). The recording methodology follows (Hromadka et al., 2008; Katz et al., 2012). Specifically, a glass micropipette (resistance 3-5 MΩ) was filled with cortex buffer and lowered by a PatchStar manipulator (Scientifica, UK) to a depth of 316–750 µm (median 546 µm) below the pia. While lowering the pipette, the resistance at the tip was monitored by applying current pulses. On approach to cells, negative pressure was applied to form a seal (20–1000 MΩ) with the cell membrane. The signals were amplified using Multiclamp 700B amplifier (Molecular Devices, USA) and sampled at 30 kHz for storage and offline analysis.

Multi-site extracellular recordings were performed in 28 mice (**Table S2**) with either Buzsaki32 (4 shanks, 8 electrode sites per shank) silicon probes or A1×32-Edge-5mm-20-177 (‘Edge32’) probes (both from NeuroNexus Technologies, USA). The probe was lowered to a depth of 535–845 µm (median 615 µm) and 2320–3000 µm (median 2675 µm) for V1 and LGN recordings, respectively. We identified the electrode sites in the LGN by testing for visual responses with 1-s reversing checkboard stimuli presented on liquid-crystal display (LCD) monitors in front of the contralateral eye (confirmed post-hoc by first spike-sorting with KiloSort (Pachitariu et al., 2016) and Phy (Rossant et al., 2016), and then testing the units for rhythmic responses to reversing checkerboard stimuli). The probe was advanced until robust visual responses were seen. We recorded simultaneously from the LGN and V1 in five sessions in four mice (**Table S2**).

Signals were amplified, sampled at 30 kHz, and stored for offline analysis using the Cerebus data acquisition system (Blackrock Microsystems, USA).

### S1.5 Optogenetic stimuli and flash visual stimuli

Optogenetic stimulation was applied to the cortex while the mouse was facing a blank gray screen that was shown on three LCD monitors, covering a field of view of roughly 120° x 60° extending in front and to the right of the animal.

To perform dichromatic optogenetics, we used a LightHub system (Omicron-Laserage Laserprodukte GmbH, Germany), incorporating a blue laser (445 nm, Omicron LuxX 445-100) and a green laser (561 nm, Omicron DPSS laser). The laser system was coupled to a 400-µm optic fiber with a collimator (F280FC-A; Thorlabs, USA) and a lens (f = 50 mm; Thorlabs, USA), which was positioned a few centimeters above the pial surface and centered on the V1 recording site. The laser pulses were at the maximum power possible (100 and 150 mW for the blue and green laser, respectively). Optogenetic stimuli included single laser pulses of various durations (0.5–4 ms) and paired pulses (0.5– 2 ms) to stimulate E and I neurons in all four possible orders with interpulse intervals (IPIs) ranging from 5 to 800 ms (**Table S3**). Optogenetic stimuli were applied in trial blocks: each block contained a single repeat of each stimulus, arranged in a random order that differed from block to block. There were 10–68 blocks per recording session; many trial blocks contained only single pulses (**Table S2**). Between each pulse (or each pair of pulses for paired-pulse stimuli) there was an interval of 1.6–5 s, chosen at random.

For the LGN and simultaneous LGN–V1 recordings, we successively positioned the laser above different parts of exposed V1 to ensure that we triggered the part of V1 directly connected to the recorded LGN neurons. The optimal laser position was identified by the magnitude of the initial LGN response (multi-unit activity) to the laser pulse. The larger the initial multi-unit activity, the more likely that the stimulated V1 region was directly connected to the recorded LGN neurons. Recording sessions in which no initial LGN response was higher than five times the baseline activity were discarded (two sessions). In some recording sessions, the LGN neurons showed visual responses to the laser pulse, possibly because the light hit the back of the retina through the brain. These sessions were identified by response latency and discarded from further analysis.

Flash visual stimuli were presented to wild-type mice. These stimuli were generated by briefly flashing an LED (1–4 ms; 470 nm, power ∼5 mW/cm^2^; ThorLabs, USA) to the contralateral eye of a mouse in darkness. Similar to optogenetic stimuli, there was 2.1–2.5 s gap between flashes. (In one session, the gap was between 15–30 s.) Video monitoring showed that pupil size decreased immediately after light exposure and recovered back to its baseline size before the onset of the next light flash.

### S1.6 Spike detection

To estimate the total firing rates of excitatory and fast-spiking inhibitory neuronal populations, we developed a “clusterless” algorithm to distinguish wide spikes, typical of excitatory neurons, from narrow spikes, typical of cortical parvalbumin-expressing interneurons (Bartho et al., 2004), without having to spike-sort.

The method classifies each spike individually as narrow or wide. The waveform of a single raw spike is too noisy to measure its waveform characteristics accurately, so we applied a local smoothing method that denoised the unfiltered waveform by averaging together the waveforms of multiple similar spikes, without requiring explicit clustering (**Fig. S2A**). For each target spike, we applied locality-sensitive hashing^1^ to find (up to) 100 spikes of similar waveform, assessed by Euclidean distance on the PCA features produced by SpikeDetekt (Rossant et al., 2016), that therefore have high probability of being fired from the same neuron. We then averaged these spikes to estimate a smooth waveform for the target spike.

Multiple quality control criteria were used to reject spikes whose waveforms could not be reliably identified using this method. First, badly smoothed spikes were identified using a normalized fitting error: the RMS error between the filtered waveform of the target spike and the smoothed waveform, divided by the peak-to-trough amplitude of the smoothed waveform. Spikes were discarded if the normalized fitting error exceeded a threshold of 0.45. Second, spikes whose smoothed waveform had a small amplitude (maximum amplitude of the filtered waveform, amp, < 25 µV) were rejected as noise. Third, target spikes whose distance to their 5^th^ nearest neighbor exceeded 0.305 (after normalization by peak-to-trough amplitude of the smoothed spike) were rejected. Finally, spikes were rejected if their normalized distance to their 1^st^ nearest neighbor exceeded 0.0305 and the normalized fitting error exceeded 0.305. These criteria and parameters were chosen after manual inspection of all recordings, and by cross-checking on a subset of recordings in which the method produced similar results to traditional spike-sorting methods. For the latter analysis, we compared the PSTHs obtained by our locality-sensitive hashing method to PSTHs obtained after combining narrow- and wide-spiking clusters obtained by traditional spike-sorting methods into E and I population rates.

To ensure that movement artifacts and photoelectric artifacts caused by optogenetic stimulation are not considered as genuine spikes, spike events whose waveforms were similar across recording sites (variability < 2.53 µV^2^ and variability < 5.66 µV^2^ if variability/amp < 0.05 µV) were rejected.

Spikes passing these quality-control criteria were then classified as wide or narrow using four waveform characteristics (**Fig. S2B)**: (1) spike duration, measured between the trough and the following peak of the smoothed unfiltered waveform, (2) full width at 2/3 maximum (FW3M), and (3) early and (4) late spike gradients, measured at 0.07 and 0.50 ms after the waveform trough. Spikes were classified as narrow or wide by defining two regions within the four-dimensional space defined by these four parameters. The boundaries of these regions were chosen conservatively to ensure a low false positive rate (by manual inspection and cross-check against clustered data), and spikes outside of these regions were discarded from further analysis (see **Fig. S2C** for an example). The boundaries for spike duration and FW3M were chosen manually for each recording sessions to maximize the number of detected spikes per session, consistent with these conservative criteria:

|  | Wide spikes 'excitatory' | Narrow spikes 'fast spiking inhibitory' |
| --- | --- | --- |
| **Spike duration (ms)** | 0.47–0.83 | 0.13–0.33 |
| **FW3M (ms)** | 0.18–0.30; 0.18–0.33; 0.15–0.30 | 0.08–0.18; 0.07–0.20; 0.10–0.18; 0.10–0.22 |
| **Early gradient (μV/ms)** | < 1.16 | > 1.16 |
| **Late gradient (μV/ms)** | > 0 | < 0 |

This method allowed us to classify as narrow or wide ∼40 % of all spike events produced by SpikeDetekt. The remainder were excluded from further analysis.

For LGN recordings, a distinction between narrow and wide spikes was not made, and multi-unit activity was estimated by pooling the spikes of all spike-sorted units with rhythmic responses to reversing checkerboard visual stimuli.

### S1.7 Data analysis: peristimulus time histograms

Peristimulus time histograms (PSTHs) were computed using a bin size of 1/30 ms (the hardware’s sampling frequency) and smoothed with Hamming windows of appropriate lengths. For juxtacellular recordings, spikes from all neurons were summed into a virtual population and smoothed with either 1- or 40-ms Hamming window (**Fig. 1E**); while PSTHs of single neurons (used to extract recovery time) were smoothed by a 100-ms Hamming window because of much lower spike counts (**Fig. 1D**). For extracellular recordings, we computed smoothed PSTHs of the E and I population responses for every stimulus condition using a 40-ms Hamming window and averaging across trials of each stimulus condition. Finally, we normalized all smoothed, trial-averaged PSTHs by dividing by each recording’s mean baseline activity (activity between 500 and 100 ms before the first laser pulse, across stimulus conditions) so that the pre-stimulus baseline rates of E and I cells were both 1.

We characterized the impulse-response dynamics by two numbers (**Fig. 2D**, inset): the recovery time (defined as the time that the E population response, assessed by the PSTH calculated as described previously, recovered to 50% of its baseline activity) and the rebound amplitude (defined as the peak E population response post-suppression). For single-pulse experiments, we sometimes applied optogenetic pulses (or visual flashes) of various durations. For such sessions, we averaged the data (i.e., the recovery time and the rebound amplitude) across all pulse durations to obtain a single value per session for statistics and display purposes in Fig. 2D.

### S1.8 Model fitting

Models were fit and tested using cross-validation (see S1.10). To find the models’ parameters from the training-set data, we optimized a loss function measuring the similarity between the models’ predicted responses and the PSTHs computed from the training-set data. The models’ differential equations were integrated using Euler’s method, and the loss function was optimized using the simplex method (‘fminsearch’ in MATLAB).

The loss function added together error functions measuring the mismatch between the model-predicted and experimentally measured E and I population rates for each stimulus type (i.e., single pulses, paired pulses, and flash visual stimuli) in the training set. The loss function *L* was defined as:

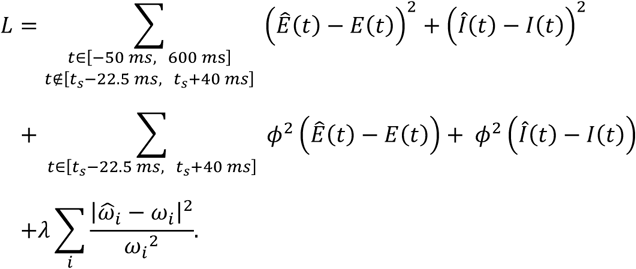

Here, *E*(*t*) and *I*(*t*) are the experimentally measured PSTHs of the E and I populations, after smoothing with a 40-ms Hamming window, trial-averaging, and baseline-division. *Ê* (*t*) and *Î* (*t*) are the corresponding simulated data, smoothed with the same 40-ms Hamming window as the experimental data to allow fit-errors to be estimated by simple subtraction. The onset of a pulse is denoted by *t*_*s*_; the first pulse is always at time 0, but there can be a second pulse for paired-pulse experiments. The first summand in the above equation is the sum of squared errors over all time points in the indicated range (which starts before time 0 because of non-causal smoothing), excluding a window around the pulse period when spike detection was compromised, at a bin resolution of 1/30 ms (the hardware’s sampling frequency). The second summand is an error term for the pulse period (colored in ivory in **Fig. 4B–E, 5B–E, 7B–E**). Because experimentally measured firing rates around the pulse period are underestimated, but are unlikely to be overestimated, errors during this period were only counted if the simulated rate was smaller than the experimental measurement. This was achieved by passing the errors through a squared rectified linear function: *ϕ*^2^(*x*) = (max(*x*, 0))^2^.

The final summand represents a regularization term, with 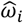 the estimated parameters, *ω*_*i*_ a manually chosen prior. When fitting dynamical models to data, there are often multiple sets of parameter values that can give an equally good fit (Prinz et al., 2004). Our goal in the current study was not to find specific parameter values, but rather to determine which classes of models can accurately fit the data. We therefore used the strategy of annealing a prior to zero, which allows the model fit to avoid local minima, while having no effect on the precise final value. The regularization penalty strength *λ* started at 100 and was gradually annealed to 0 in successive rounds of optimization (step size 10). Each round ran until convergence criteria TolX = 1e-3 and TolFun = 1e-4 were reached. TolFun is the termination tolerance for the value of the loss function; TolX is the termination tolerance for the parameters.

For one recording session, because of the low baseline activity of the I population (<15 spikes/s), we reduced the contribution from the *I* term by a fifth.

### S1.9 Model equations

The differential equations used for each model were given in the main text and figures. The external inputs *X*_*E*_ and *X*_*I*_ are functions of time. Because ChR2 has rapid kinetics, we could model the time course of *X*_*E*_ by a boxcar function across the duration of the laser pulse, with amplitude multiplier the only parameter being a variable. The time course of *X*_*I*_, however, was modeled by an *α*-function *te*^−*t*/*τ α*^ to account for the slower kinetics of C1V1_T/T_, with *τ*_*α*_ another optimized parameter.

The WC model has 11 free parameters (*τ*_*E*_, *τ*_*I*_, g_*EE*_, *g*_*IE*_, *g*_*EI*_, *g*_*II*_, *X*_*E*_, *X*_*I*_, *τ*_*α*_, *E*_max_, and *I*_max_). *C*_*E*_ and *C*_*I*_ were inferred analytically: given the other parameters, they were uniquely determined as the only possibilities giving baseline rates of 1 for both E and I populations. The WCS model has an additional 3 tunable parameters (*τ*_*S*_, *g*_*ES*_, *g*_*IS*_), while the CT model has a further 11 free parameters (*τ*_*B*_, *τ*_*R*_, *g*_*EL*_, *g*_*IL*_, *g*_*LE*_, *g*_*LB*_, *g*_*LT*_, *g*_*RE*_, *g*_*RL*_, *C*_*R*_, and *L*_max_).

The pre-pulse initial conditions for all simulations were *E*_0_ = *I*_0_ = *S*_0_ = *L*_0_ = 1 and *R*_0_ = *B*_0_ = *T*_0_ = 0, which represents a stable fixed point of the dynamical system. To place a hard constraint on parameters (e.g., to insist connectivity terms be positive), the loss function was set to infinity if any of the following conditions were not satisfied during the optimization procedure:

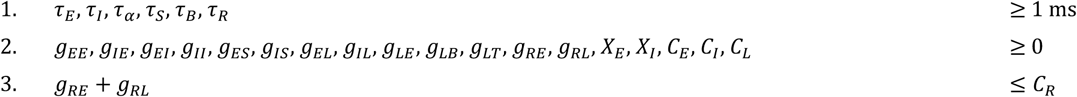

To account for session-to-session variability, we fit separately models to each recording session and tested them with cross-validation (S1.10); within each session, we fit models separately to single-pulse and paired-pulse data.

### S1.10 Validation methods

We assessed each model’s performance using cross-validation adapted to the three experimental paradigms we used: single-pulse data (11 datasets with responses to both E pulse and I pulse), paired-pulse data (9 datasets), and flash visual stimuli (6 datasets). Different procedures were applied as the latter two datasets allow a more stringent form of cross-validation (out-of-sample testing) than the single-pulse dataset.

To assess the models’ predicted responses to single pulses (**Fig. S5A**), we first randomly divided all trials into 3 folds, then assessed the model’s performance by 3-fold cross-validation. For a recording session that has *k* stimulus blocks (i.e., *k* trials of each pulse type), we divided the trials so that each fold has int(*k/3*) randomly chosen trials of single E and I pulse. Then for the first iteration, we trained the model on the PSTHs averaged across the trials in the second and third folds, and assessed its performance on the PSTHs averaged across the trials in the first fold, and so on.

To assess the models’ predicted responses to paired pulses, we used a *k*-fold out-of-sample validation method to verify that models trained on some interpulse intervals (IPIs) could generalize to other IPIs (**Fig. S5B**). We first randomly divided the *N* IPIs in a given recording session into *k* folds so that each fold contained int(*N/k*) IPIs. Since different recording sessions used different set of IPIs, *k* could be either 2 or 3 (see **Table S3** for a detailed breakdown). For each of these folds, we trained the model on the other *k-1* folds, and assessed the model’s performance on the held-out data. These steps were repeated for each of the *k* folds as a test set. Note that for every IPI, there are four possible paired-pulse combinations (I→E, E→I, E→E, and I→I stimulation), meaning that for every iteration, the model was trained on 4(*k*-1)*N*/*k* stimulus conditions, and its performance assessed on the remaining 4*N*/*k* conditions.

To assess the models’ predicted responses to flash visual stimuli, we used a leave-one-out method for out-of-sample testing (**Fig. S5C**). There are four stimulus conditions (i.e., 1-, 2-, 3-, and 4-ms light flash) in each recording session. For each stimulus condition, we trained the model on the other three conditions, and tested it on the remaining condition. These steps were repeated for each of the four conditions as a test set. The ability of the model to generalize across these conditions shows that the duration of the light flash has only a minor effect on the impulse-response dynamics (see also **Fig. S3**).

### S1.11 Modification to the models and the loss function to fit responses to flash visual stimuli

Because signals from retina arrive to LGN rather than to cortex, we modified the CT model when fitting them to flash visual stimuli. The modified CT model is:

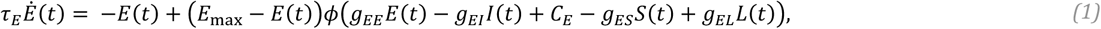

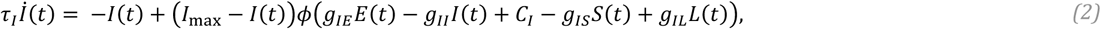

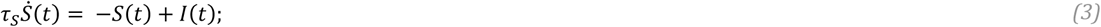

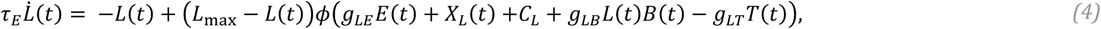

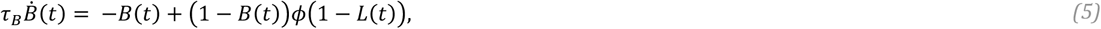

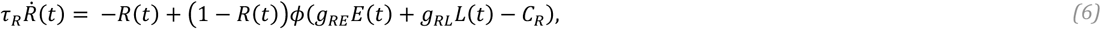

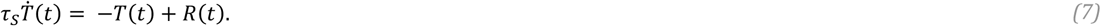

Here, *X*_*L*_ is the external input to LGN due to the light flash. Its time course is modeled as a constant function starting 20 ms after the onset of the light flash (to account for retinothalamic latency) and lasting for the duration of the light flash (i.e., *X*_*L*_(*t*) = constant for 20 ms ≤ *t* < 20 ms + light-flash duration; *X*_*L*_ = 0 otherwise).

The two cortical models (the WC and WCS models) were not modified, except for a 25 ms delay relative to the onset of the light flash (i.e., *X*_*E*_ ≠ 0 and *X*_*I*_ ≠ 0 for 25 ms ≤ *t* < 25 ms + light-flash duration; *X*_*E*_, *X*_*I*_ = 0 otherwise). The additional 5 ms delay accounts for the delay from LGN to V1.

In addition, for all three models, we extended the exclusion zone (where underestimation of population rates is expected) to 75 ms after the light-flash onset to account for the delay in visual responses and that visual responses tended to be slightly broader than responses to optogenetic stimulation.

## Footnotes

1 The locality-sensitive hashing code was taken from https://ttic.uchicago.edu/~gregory/download.html

